# Mechanism and site of action of big dynorphin on ASIC1a

**DOI:** 10.1101/816264

**Authors:** C.B. Borg, N. Braun, S.A. Heusser, Y. Bay, D. Weis, I. Galleano, C. Lund, W. Tian, L.M. Haugaard-Kedström, E.P. Bennett, T. Lynagh, K. Strømgaard, J. Andersen, S.A. Pless

## Abstract

Acid-sensing ion channels (ASICs) are proton-gated cation channels that contribute to neurotransmission, as well as initiation of pain and neuronal death following ischemic stroke. As such, there is a great interest in understanding the *in vivo* regulation of ASICs, especially by endogenous neuropeptides that potently modulate ASICs. The most potent endogenous ASIC modulator known to date is the opioid neuropeptide big dynorphin (BigDyn). BigDyn is upregulated in chronic pain and increases ASIC-mediated neuronal death during acidosis. Understanding the mechanism and site of action of BigDyn on ASICs could thus enable the rational design of compounds potentially useful in the treatment of pain and ischemic stroke. To this end, we employ a combination of electrophysiology, voltage-clamp fluorometry, synthetic BigDyn analogs and non-canonical amino acid-mediated photocrosslinking. We demonstrate that BigDyn binding results in an ASIC1a closed resting conformation that is distinct from open and desensitized states induced by protons. Using alanine-substituted BigDyn analogs, we find that the BigDyn modulation of ASIC1a is mediated through electrostatic interactions of basic amino acids in the BigDyn N-terminus. Furthermore, neutralizing acidic amino acids in the ASIC1a extracellular domain reduces BigDyn effects, suggesting a binding site at the acidic pocket. This is confirmed by photocrosslinking using the non-canonical amino acid azido-phenylalanine. Overall, our data define the mechanism of how BigDyn modulates ASIC1a, identify the acidic pocket as the binding site for BigDyn and thus highlight this cavity as an important site for the development of ASIC-targeting therapeutics.

**Significance Statement:** Neuropeptides such as big dynorphin (BigDyn) play important roles in the slow modulation of fast neurotransmission, which is mediated by membrane-embedded receptors. In fact, BigDyn is the most potent known endogenous modulator of one such receptor, the acid-sensing ion channel (ASIC), but the mode of action remains unknown. In this work, we employ a broad array of technologies to unravel the details of where big dynorphin binds to ASIC and how it modulates its activity. As both BigDyn and ASIC are implicated in pain pathways, this work might pave the way towards future analgesics.

## Introduction

Neuropeptides are a diverse class of signalling molecules that are involved in a wide variety of physiological functions, including the modulation of neurotransmission (1, 2). The neuropeptide subclass of dynorphins is best known for its modulation of the G protein-coupled opioid receptors (3–5), through which they mediate spinal analgesia. However, it is increasingly recognised that these highly basic peptides also modulate the activity of ionotropic receptors, such as tetrameric glutamate receptors (N-methyl-D-aspartate (NMDA) subtype) and trimeric acid-sensing ion channels (ASICs) (6–10). The latter interaction is of particular interest, as ASICs have emerged as mediators of both pain and stroke and thus represent potential targets in the treatment of these diseases (11–17). In fact, big dynorphin (BigDyn) is the most potent endogenous ASIC modulator described to date. The neuropeptide was found to rescue proton-gated currents after exposure to steady-state desensitization-inducing conditions in homomeric ASIC1a and heteromeric ASIC1a/2a and ASIC1a/2b channels in the nanomolar range and thereby promote acidosis-induced neuronal cell death in cultured neurons (10, 18). As ASIC1a homomers and ASIC1a-containing heteromers are the most prevalent isoforms in the nervous system, this raises the possibility that inhibitors or competitors of the ASIC1a-BigDyn interaction might prove valuable therapeutics. However, despite this potential therapeutic relevance, the underlying mechanism behind this potent modulation remains elusive.

BigDyn (32 aa) and its two smaller peptide fragments, dynorphin A (DynA, 17 aa) and dynorphin B (DynB, 13 aa), are cleavage products of the prodynorphin precursor peptide (4, 5, 19). DynA is evolutionarily highly conserved among mammals and amphibians and displays, along with BigDyn, nanomolar affinities for a range of related opioid receptors (4). BigDyn was reported to be ∼1000-fold more potent in its modulation of ASIC1a steady-state desensitization compared to DynA, while DynB did not modulate ASIC1a steady-state desensitization (10). This raises the possibility of a unique ASIC-selective pharmacophore.

Here, we set out to investigate the molecular determinants of the high potency modulation of ASIC1a by BigDyn and identify the binding site on ASIC1a. Interestingly, previous data on the well-studied ASIC1a-modulator spider toxin psalmotoxin-1 (PcTx1), which increases the apparent proton sensitivity and consequently enhances steady-state desensitization (20), pointed towards a possible role of the ASIC1a acidic pocket in BigDyn modulation: firstly, pre-incubation in PcTx1 has shown to prevent BigDyn modulation of ASIC1a, suggesting that the two peptides might compete for an overlapping binding site (10). Secondly, the positively charged toxin is bound at the acidic pocket in ASIC1a–PcTx1 co-crystal structures (21, 22), again raising the possibility of a similar binding site for BigDyn, which carries a net charge of +9.

In order to directly probe the molecular determinants of the high-potency modulation of ASIC1a by BigDyn, we employ a multi-angle strategy: we introduce complementary charge-neutralizing mutations in ASIC1a, as well as BigDyn and study their functional consequences on channel modulation in electrophysiology experiments. We further use using voltage-clamp fluorometry (VCF) recordings to monitor the conformational changes resulting from the ASIC1a-BigDyn interaction. Finally, the non-canonical amino acid photocrosslinker azido-phenylalanine (AzF) (23, 24) is incorporated at the acidic pocket of ASIC1a to covalently crosslink ASIC-BigDyn complexes using UV exposure. Together, our results demonstrate that BigDyn binds at the acidic pocket through charge-charge interactions and reduces the proton sensitivity of both activation and steady-state desensitization of ASIC1a, likely by inducing a distinct closed state of the channel.

## Results

### BigDyn affects pH dependence of mASIC1a and induces a unique closed state

First, we set out to test the ability of BigDyn and its proteolytic products (Figure 1A) to rescue proton-gated currents after exposure to steady-state desensitization-inducing conditions of wild-type (WT) mouse ASIC1a (mASIC1a) using two-electrode voltage clamp (TEVC) electrophysiology. Steady-state desensitization was induced using conditioning in pH 7.1, which decreased subsequent pH 5.6-induced current response to 5.7 % (95 % confidence interval (95CI): 3.8, 7.6 %) compared to that after conditioning in pH 7.4 (“Control”). However, application of 1 µM BigDyn during the pH 7.1 conditioning step largely rescued pH 5.6-induced currents, resulting in 90.1 % (95CI: 77.9, 102.3 %) of the control current response. The EC_50_ for this current rescue was 210.6 nM (95CI: 162.6, 258.6 nM) BigDyn; Figure S1A-B). By contrast, applying DynA and DynB during conditioning did not produce significant current rescue when applied at a concentration of 1 µM (at pH 7.1) (Figure 1B-C), although at a concentration of 3 µM, DynA partially rescued pH 5.6-induced current (Figure S1C). Furthermore, co-application of 1 µM DynA or Dyn1-19 (containing the Arg18 and Lys19 that link DynA and DynB in full-length BigDyn) with DynB during conditioning did not reproduce the phenotype observed with full-length BigDyn (Figure 1B-C). Next, we sought to assess if the presence of BigDyn would affect the pH dependence of activation or steady-state desensitization in WT mASIC1a. Indeed, even the presence of a sub-saturating BigDyn concentration (0.1 µM) resulted in a small right-shift in the pH_50_ of activation (from 6.80 (95CI: 6.70, 6.91) to 6.66 (95CI: 6.61, 6.70), p = 0.0172) and of steady-state desensitization (from 7.21 (95CI: 7.19, 7.24) to 7.15 (95CI: 7.12, 7.19), p = 0.0225) (Figure 1D, Table S1).

**Figure 1:**
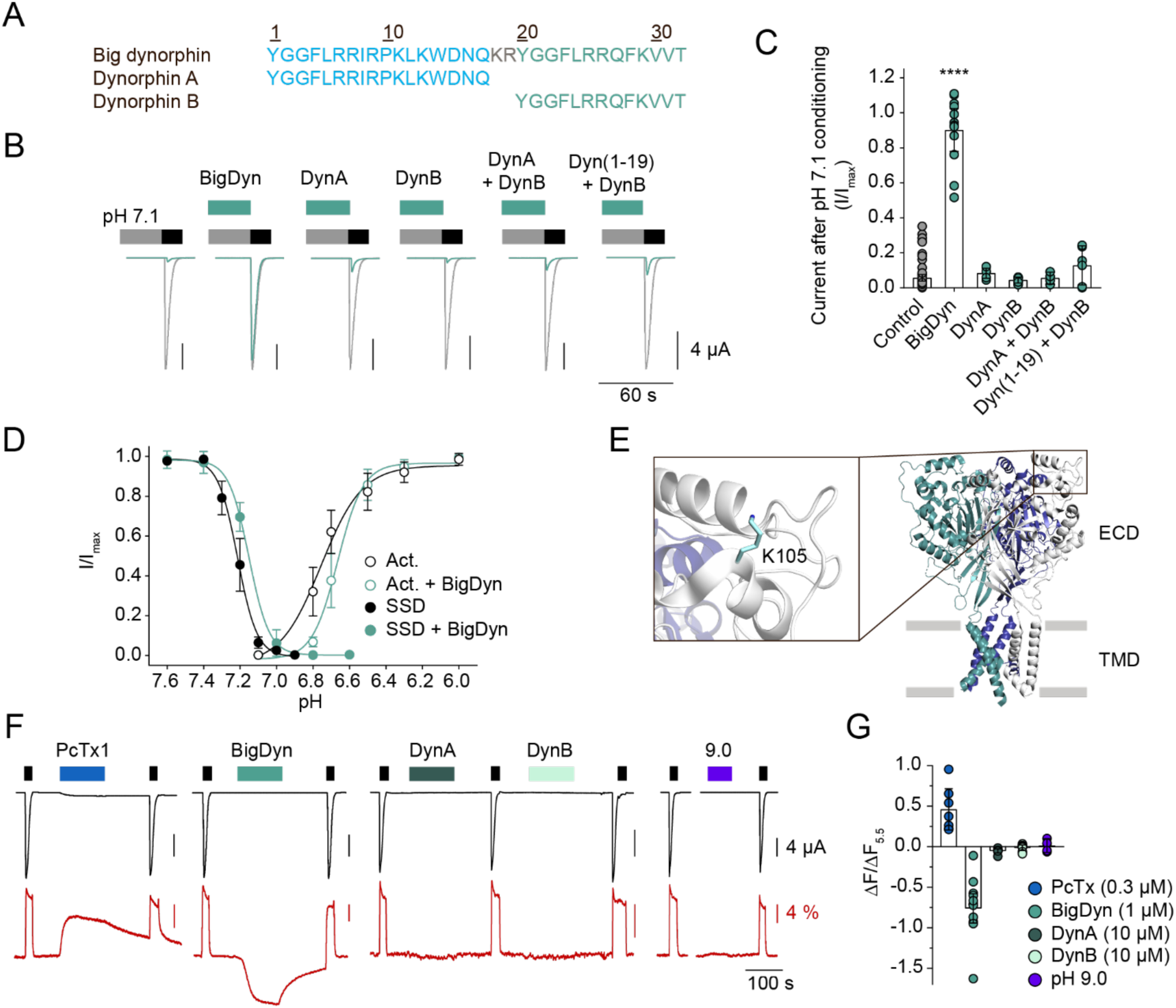
Mechanism of ASIC1a modulation by BigDyn. (A) Amino acid sequences of BigDyn, DynA and DynB. (B/C) Representative current traces (B) and averaged data (C) obtained by pH 5.6 application (black bar in (B)) at mASIC1a WT-expressing *Xenopus laevis* oocytes after preincubation in pH 7.1 (grey bar) with or without 1 µM of the indicated peptide or peptide combination (green bar); in (B) control currents (after pH 7.4 conditioning) are shown in grey for comparison. Asterisk in (C) indicates significant difference to control condition (p < 0.0001); n = 5-68). (D) Concentration-response curves for activation (Act.) and steady-state desensitization (SSD) of WT ASIC1a in the presence and absence of 0.1 µM BigDyn (n = 4-15). (E) Structure of cASIC1a (PDB: 4NTW) with individual subunits color coded and inset showing the location of Lys105. (F) Representative current (black) and fluorescence (red) traces obtained by application of pH 5.5 (black bars) at mASIC1a labeled with Alexa Fluor 488 at position 105, the indicated peptide (PcTx1, 0.3 µM: blue bar; BigDyn, 1 µM: green bar; DynA, 10 µM: dark green bar, and DynB, 10 µM: light green bar) or pH 9.0 (purple bar). (G) Averaged change in fluorescence obtained by application of PcTx1, BigDyn, DynB or pH 9.0, as shown in (F) (normalized to that obtained by application of pH 5.5) (n = 5-15). Error bars in (C), (D) and (G) represent 95CI.

The above results suggested that the potency of both proton activation and steady-state desensitization was lowered in the presence of BigDyn, but it remained unclear if this occurred through a stabilization of the canonical resting closed state (relative to open/steady-state desensitized states) or if BigDyn induced a distinct protein conformation. To distinguish between these possibilities, we turned to VCF, which allows us to directly probe ion channel conformational rearrangements using environmentally sensitive fluorescent dyes (25). In agreement with earlier reports (26), Alexa Fluor 488-labelled Lys105Cys mutants (Figure 1E) showed robust and reversible pH-induced current and fluorescence changes (Figure 1F, Table S2). The upward deflection of the fluorescence signal is likely associated with channel desensitization, as the pH response curve of the fluorescence closely matches that of steady-state desensitization (Figure S2A, Table S3). Additionally, application of 0.3 µM tarantula toxin PcTx1, reported to stabilize the desensitized state of ASIC1a by increasing the proton sensitivity (20, 27) through binding to the acidic pocket (21, 22), also induces an upward deflection in fluorescence (dequenching) without inducing current (Figure 1F-G). By contrast, application of 1 µM BigDyn did not result in a current response, but caused a downward deflection (quenching) of the fluorescence signal, in stark contrast to the upward deflection observed with increased proton concentrations and PcTx1. This quenching is dependent on the BigDyn concentration, with a half-maximal effect in the high nanomolar range (Fig. S2B), slightly higher than what was observed for its functional EC_50_ on the WT channel (850 nM (95CI: 299, 1402 nM) and 211 nM (95CI: 163, 259 nM), respectively). Importantly, no fluorescence change was observed upon application of 10 µM DynA or 10 µM DynB or increasing the pH to 9.0 (Figure 1F-G, Table S2), demonstrating that the BigDyn-induced conformation is dependent on a functionally active peptide and distinct from that elicited by low (or high) pH. Together, the data show that mASIC1a modulation by BigDyn is dependent on a continuous peptide sequence containing, at least parts of, both DynA and DynB and indicates that BigDyn might be stabilizing a closed resting conformation of ASIC1a that is distinct from that induced by protons and PcTx1.

### Contribution of positively charged BigDyn side chains to modulation of ASIC1a

An unusual feature of BigDyn is its high density of positive charge (net charge +9, Figure 2A). We therefore reasoned that the interaction with ASIC1a might be driven by electrostatic forces, i.e. by binding to negatively charged side chains on ASIC1a. To investigate the contribution of individual positively charged amino acids (Arg and Lys) of BigDyn to its ability to modulate ASIC1a steady-state desensitization, we generated BigDyn analogs with individual positively charged residues substituted for alanine (Ala). These ten BigDyn Ala-analogs were tested for their ability to rescue current at WT mASIC1a when applied at 1 µM at a conditioning pH of 7.1 (Figure 2B-C). Indeed, individual alanine substitutions of the three most N-terminally located positively charged amino acids (Arg6, Arg7 and Arg9) virtually abolished BigDyn-mediated current rescue: 9.8 % (95CI: −2.1, 21.6 %) for Arg6Ala, 5.0 % (95CI: 1.2, 8.7 %) for Arg7Ala, and 2.3 % (95CI: 1.7, 3.0 %) for Arg9Ala, respectively (Table S4).

**Figure 2:**
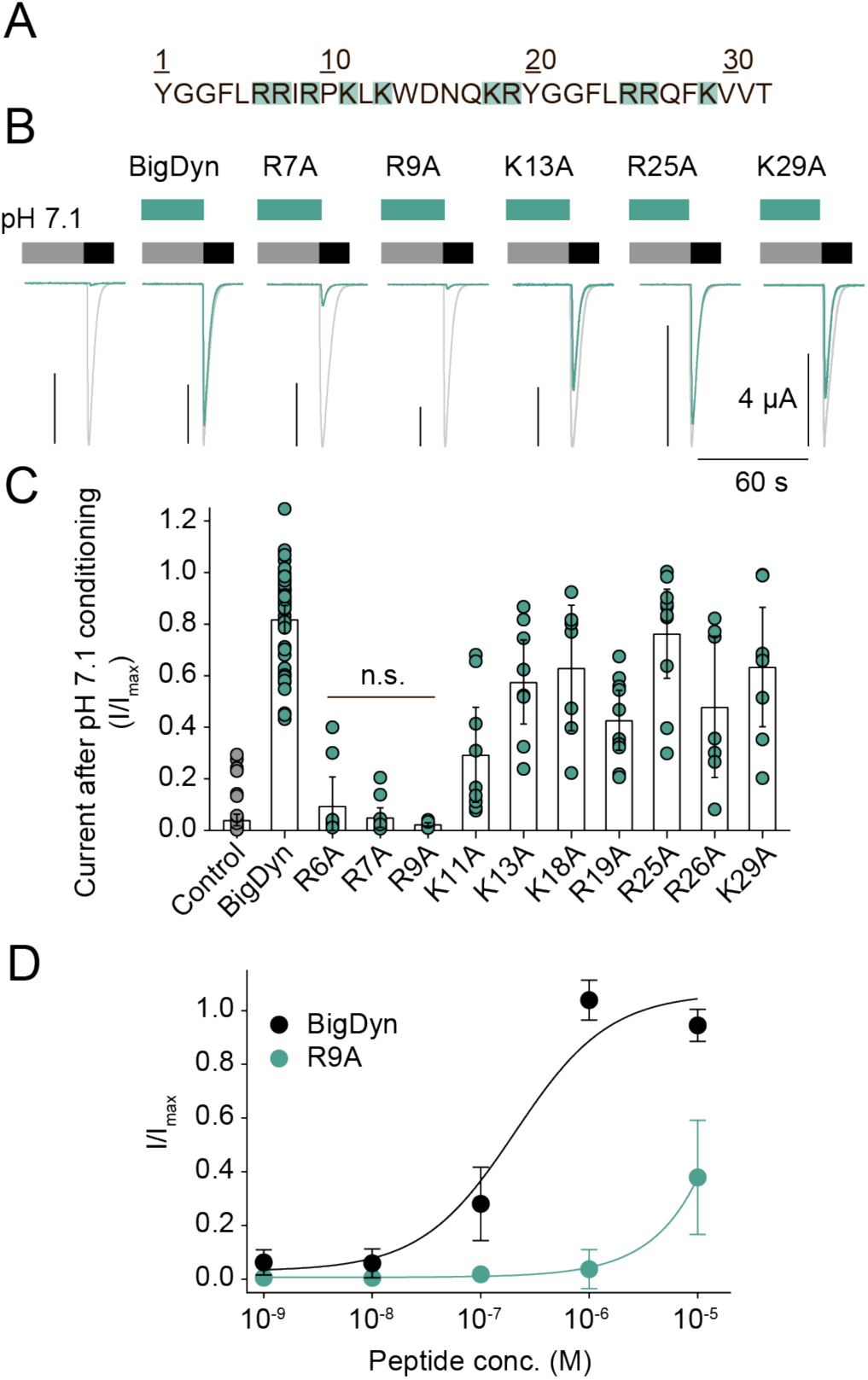
Positive charges near the BigDyn N-terminus are essential for ASIC1a modulation. (A) Sequence of BigDyn with basic side chains highlighted in green. (B/C) Representative current traces (B) and averaged data (C) obtained by pH 5.6 application (black bar in (B)) at ASIC1a WT-expressing *Xenopus laevis* oocytes after preincubation in pH 7.1 (grey bar) with or without 1 µM of the indicated BigDyn analog (green bar); control currents after pH 7.6 conditioning are shown in grey for comparison (B). (D) Concentration-response curves for WT ASIC1a modulation by BigDyn (black) and Arg9Ala (green). Error bars in (C) and (D) represent 95CI. In (C), n.s. indicates no statistically significant difference to control condition. n = 7-48 in (C) and n = 7-9 in (D).

By contrast, individual substitution of the remaining positive charges (Lys11, Lys13, Lys18, Arg19, Arg25, Arg26, and Lys29; Figure 2) or the residue situated between Arg7 and Arg9 (Ile8; Figure S3A) showed less pronounced effects on BigDyn activity (between 29.4-76.3 % recovery; Table S4). This indicates that the charge of a cluster of three closely positioned Arg in the N-terminal part of BigDyn is important for the functional modulation of mASIC1a, although other properties, such as side chain size and H-bonding ability might also play a role (Figure 2). This notion is confirmed by the finding that the EC_50_ for the modulatory effect of the Arg9Ala peptide is increased over 50-fold (Figure 2D). Consistent with the above, substituting the sole negatively charged residue (Asp15) had no effect on its ability to modulate ASIC1a (Figure S3A). Replacing Tyr1, which has been implicated in the activity of BigDyn towards the opioid receptors (19, 28, 29), or Trp14, resulted in a modest, but significant decrease in the ability to rescue currents (Figure S3A), while N-terminal truncations generally had more pronounced effects (Figure S3B, Table S5).

Together, the data indicate that the BigDyn-mediated effects on ASIC1a are primarily driven by a cluster of three positively charged side chains in the N-terminal part of the neuropeptide.

### Negative charges within the acidic pocket are crucial for BigDyn modulation of ASIC1a

To complement our findings with BigDyn, we next wanted to identify its binding site on ASIC1a. As we found the basic charge of BigDyn to be a crucial determinant of the interaction, we set out to mutate negatively charged side chains in the mASIC1a extracellular domain (ECD). Specifically, we focused our attention on a cavity in the ASIC1a ECD denoted the acidic pocket, which contains a high density of negatively charged residues (30). The acidic pocket also serves as the binding site of PcTx1 (21, 22, 31), which has been suggested to compete with BigDyn for ASIC1a modulation (10). We introduced charge-neutralizing amino acid substitutions to individual acidic side chains (Asp to Asn and Glu to Gln substitutions) at eight positions in and around the acidic pocket (Figure 3A). As ASIC1a mutations, especially around the acidic pocket, often lead to a change in pH sensitivity, the pH dependence of steady-state desensitization for each of the mutant mASIC1a constructs was determined in order to identify the appropriate steady-state desensitization conditioning pH for each mutant (Figure S4, Table S6). Next, we tested the mASIC1a variants for their sensitivity towards 1 µM BigDyn, applied during steady-state desensitization conditioning. However, all tested single mutants retained significant pH 5.6-induced current rescue (Figure 3B-C, Table S7), indicating that single charge-neutralizing amino acid substitutions in or near the acidic pocket alone are not enough to abolish the effect of BigDyn.

**Figure 3:**
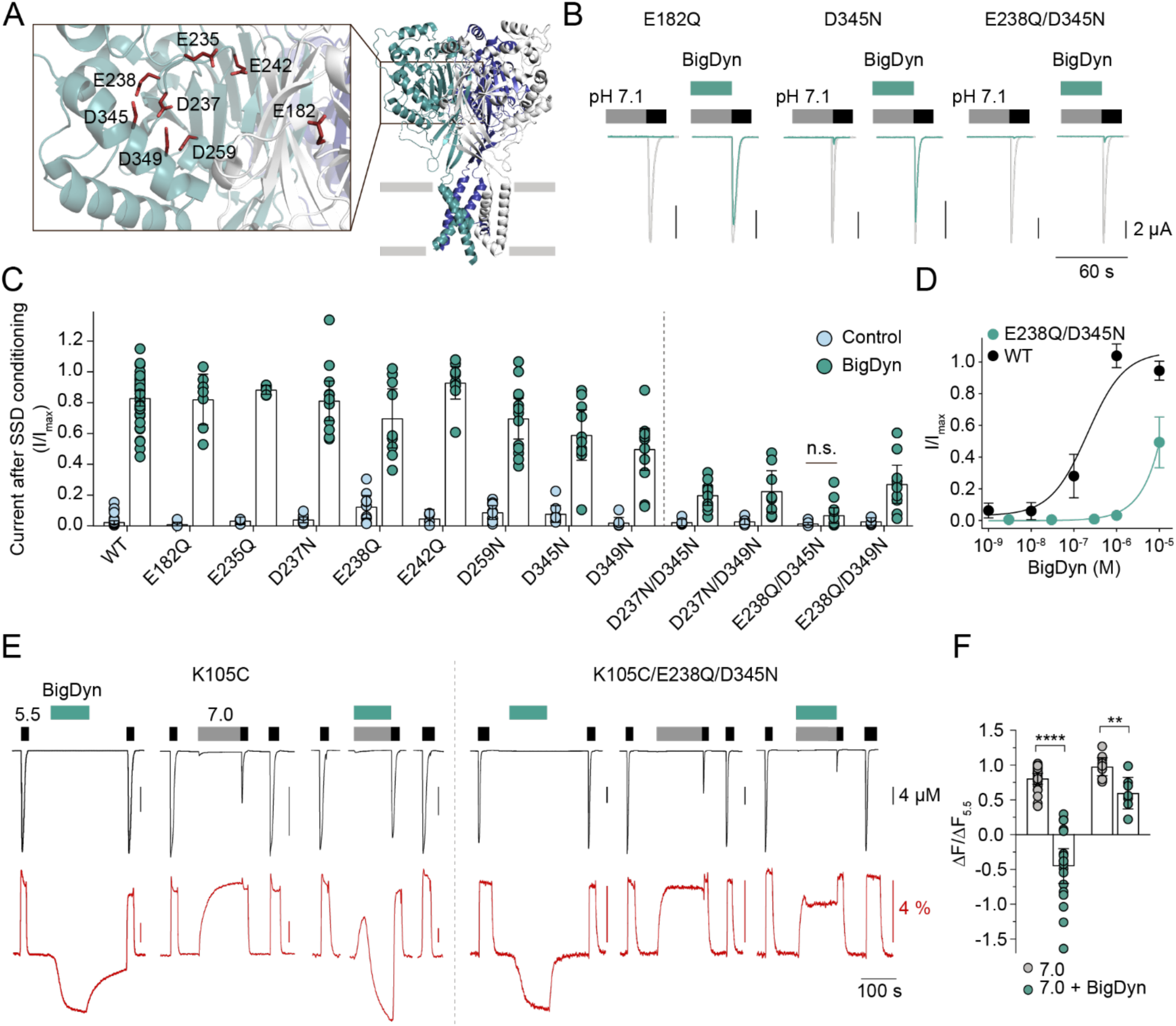
Double charge-neutralizing mutations around the ASIC1a acidic pocket reduce BigDyn modulation. (A) Structure of cASIC1a (PDB: 4NTW) with individual subunits color coded and inset showing the location of acidic side chains mutated to Gln or Asn. (B/C) Representative current traces (B) and averaged data (C) obtained by pH 5.6 application (black bar in (B)) after preincubation in steady-state desensitization (SSD)-inducing pH (grey bar; see Fig S4 and Table S6 for details) with or without (Control) 1 µM BigDyn (green bar) at *Xenopus laevis* oocytes expressing the indicated ASIC1a construct; in (B) control currents after pH 7.6 conditioning are shown in grey for comparison (n = 4-45). n.s. indicates no statistically significant difference to control condition. (D) Concentration-response curves for BigDyn-mediated modulation of WT (black) and Glu238Gln/Asp345Asn mutant (green) ASIC1a (n= 7-9). (E) Representative current (black) and fluorescence (red) traces from channels labeled with Alexa Fluor 488 at position 105 obtained by application of pH 5.5 (black bar), 7.0 (grey bar) or 1 µM BigDyn (green bar) at mASIC1a Lys105Cys (left) and Lys105Cys/Glu238Gln/Asp345Asn (right). (F) Averaged change in fluorescence at pH 7.0 with and without 1 µM BigDyn as obtained in (E), normalized to that obtained by application of pH 5.5; n = 7–22, unpaired t-test **p < 0.01; ****p < 0.0001. Error bars in (C), (D) and (F) represent 95CI.

We therefore sought to investigate whether combined mutation of two acidic residues in the acidic pocket would abolish BigDyn modulation. Four acidic residues are found in close proximity on the upper thumb and finger domains (Asp237, Glu238, Asp345, and Asp349; Figure 3A) and have been suggested to interact as two carboxyl-carboxylate pairs (Asp237-Asp349 and Glu238-Asp345) (30). We thus generated mASIC1a variants carrying a combination of two charge-neutralizing Asp to Asn and/or Glu to Gln mutations of these four residues and measured their pH dependence of steady-state desensitization (Figure S4 and Table S6). The double mutant mASIC1a constructs were then tested for their sensitivity to BigDyn modulation using the same protocol as for the single mutations, except that Asp237Asn/Asp345Asn was activated at pH 4.0 in order to reach saturating currents. All four tested acidic pocket double mutants showed a drastically decreased sensitivity to 1 µM BigDyn compared to WT mASIC1a (Figure 3B-C and Table S7). The biggest decrease in rescue of pH 5.6-induced currents was observed for Glu238Gln/Asp345Asn ASIC1a. For this construct, application of 1 µM BigDyn during steady-state desensitization conditioning resulted in 6.8 % (95CI: 1.7, 11.8 %) current compared to pH 5.6-induced currents after pH 7.6 conditioning, which was comparable to the control response observed in absence of BigDyn (Figure 3B-C). Additionally, the Glu238Gln/Asp345Asn double mutant showed a roughly 50-fold increase in the EC_50_ for the modulatory effect of BigDyn (Figure 3D). As the Hill slope of ASIC1a steady-state desensitization is unusually steep, we sought to confirm that the loss of the BigDyn effect observed for the double mutants was not due to a shift in pH sensitivity that rendered the channels overall unresponsive. As detailed in Figure S5A and Table S7, this was not the case, although the Asp237Asn/Asp349Asn mutant showed a moderately reduced effect when a different conditioning pH was employed. Additionally, we combined the Glu238Gln/Asp345Asn double mutant with the Lys105Cys mutation (used for VCF experiments) resulting in a variant that showed robust pH-induced fluorescence changes in VCF experiments. Similar to the Glu238Gln/Asp345Asn mutation alone, but contrary to the Lys105Cys single mutant, the Lys105Cys/Glu238Gln/Asp345Asn triple mutant was functionally insensitive to 1 µM BigDyn (Figure 3E, current trace). However, the application of BigDyn at pH 7.6 led to a downward deflection of the fluorescent signal to a similar extent as observed in the Lys105Cys variant (−81.7 % (95CI: −27.6 −135.9 %) and −76.0 % (95CI: −58.1, −94.0 %). When conditioning the Lys105Cys variant at pH 7.0 together with BigDyn, we observed a fluorescent signal that likely portrays dequenching due to conformational changes induced by pH 7.0 in the first phase, followed by gradually dominating quenching of the signal as a result of BigDyn binding in the later phase (Figure 3E, left). The triple mutant, however, presents a distinct fluorescent profile that is dominated by the pH 7.0 upward deflection and no net quenching despite a small BigDyn-dependent reduction in fluorescence intensity (Figure 3E–G). Together, the results indicate that BigDyn still binds to the triple mutant, but with a decreased ability to prevent steady-state desensitization.

Next, we wanted to ascertain that the loss of BigDyn modulation is indeed specific to double charge-neutralizing mutations in or near the acidic pocket. We thus tested two additional double charge-neutralizing mutations at positions outside the acidic pocket, Asp253Asn/Glu245Gln and Glu373Gln/Glu411Gln (Figure S5B-C). Both double mutants retained full sensitivity towards 1 µM BigDyn, indicating that the effect of double charge-neutralizing mutants was specific to the acidic pocket.

In principle, it is possible that the observed loss of current rescue with the double charge-neutralizing mutations in or near the acidic pocket originates from a changed desensitization profile. However, this is highly unlikely, given that there was no correlation between the extent of change in steady-state desensitization and current rescue and mutants with WT-like or even left-shifted steady-state desensitization curves showed a drastic loss of current rescue (e.g. Glu238Gln/Asp345Asn and Glu238Gln/Asp349Asn). Overall, the data therefore strongly imply that the acidic pocket serves as an interaction site for BigDyn.

### Photocrosslinking confirms an interaction site in the acidic pocket

To further validate the acidic pocket as the BigDyn binding site, we turned to UV-induced photocrosslinking using the photoreactive side chain of the non-canonical amino acid (ncAA) azido-phenylalanine (AzF) incorporated at different positions in ASIC1a (Figure 4A-B) (32–34). Incorporating AzF at positions lining the ASIC1a-BigDyn interaction interface should allow covalent trapping of the complex and enable subsequent visualization using western blotting. To this end, we removed endogenous ASIC1a from HEK 293T cells by CRISPR/Cas9 (Figure S6A) and expressed hASIC1a variants carrying AzF at Glu177, Thr236, Thr239, Lys 343, Glu344, Asp351, Glu355, Lys356 and Asp357 respectively, using the non-sense suppression methodology (Figure 4A-B) (35). Efficient ncAA incorporation was confirmed by comparing the amounts of protein obtained from cells grown in presence compared to cells grown in the absence of AzF (Figure S6B–C). Notably, while steady-state desensitization of hASIC1a is shifted compared to that of mASIC1a (hASIC1a pH_50_ steady-state desensitization: 7.07 (95CI: 7.03, 7.11) *vs.* mASIC1a pH_50_ steady-state desensitization 7.24 (95CI: 7.23, 7.25)) (36), the modulatory effect of BigDyn on the human ortholog is virtually identical (Figure S6D–F). As shown in Figure 4C, covalently crosslinked BigDyn was detected in samples containing AzF in and around the acidic pocket only when they were exposed to UV light, but not in control samples processed in absence of UV light or in UV-exposed WT hASIC1a, or when AzF was incorporated in the lower parts of the ECD, i.e. away from the acidic pocket (Phe69, Tyr71, Val80, Asp253, Trp287, Glu413). In summary, incorporation of the ncAA AzF in combination with UV-induced photocrosslinking confirmed that the BigDyn binding site is located at the ASIC1a acidic pocket.

**Figure 4:**
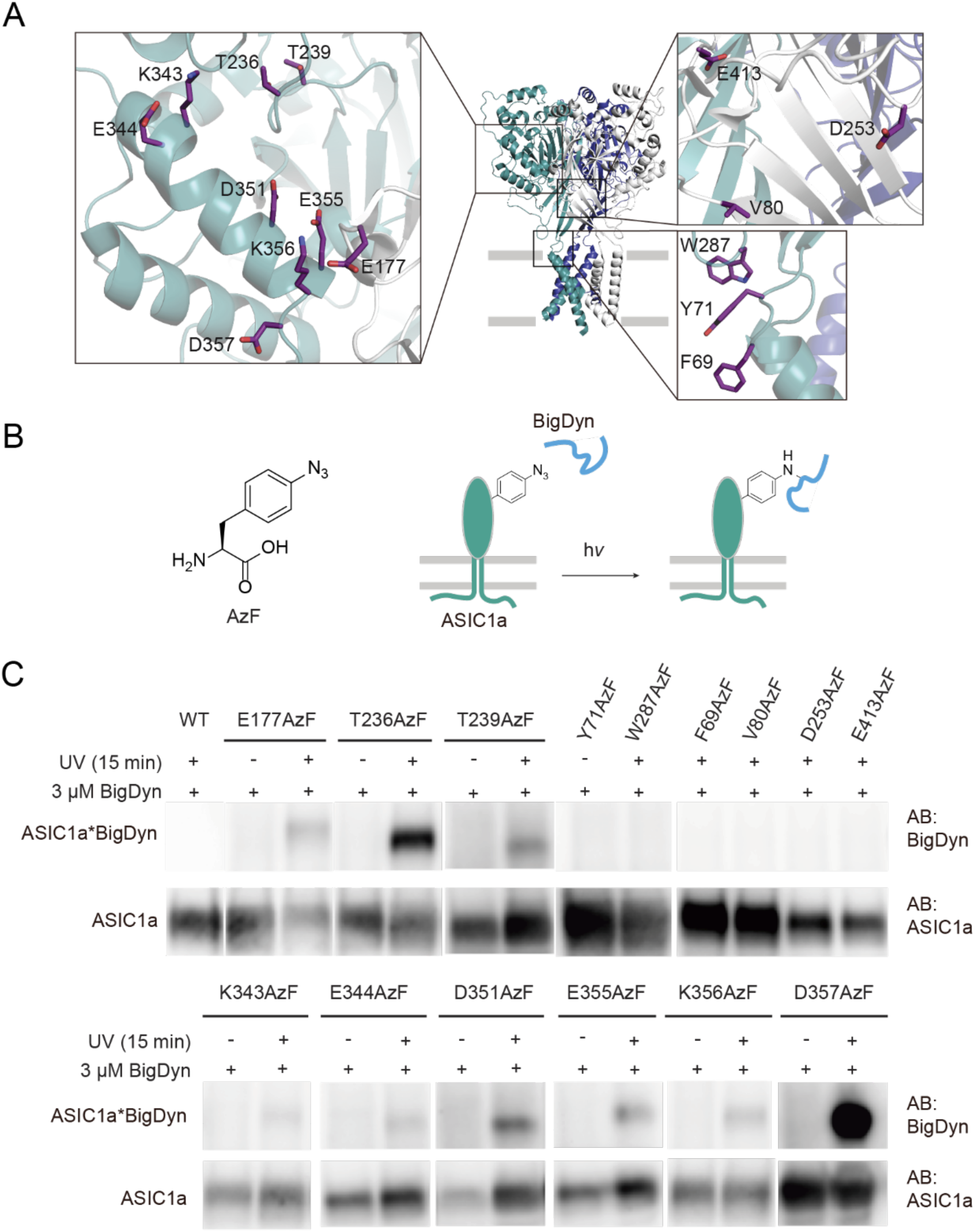
Photocrosslinking confirms the BigDyn interaction site at the acidic pocket. (A) Structure of cASIC1a (PDB: 4NTW) with individual subunits color coded and insets highlighting side chains replaced by 4-Azido-L-phenylalanine (AzF) in the acidic pocket (left inset) and lower extracellular domain (right insets). (B) Structure of 4-Azido-L-phenylalanine (AzF) and schematic workflow for crosslinking to BigDyn. HEK 293T cells expressing AzF-containing ASIC1a variants are incubated with 3 µM BigDyn and exposed to UV light for 15 min to form covalent ASIC1a-BigDyn complexes. The complex is purified via a C-terminal 1D4 tag on ASIC1a and visualized via western blotting. (C) Western blot of purified hASIC1a WT and variants carrying AzF in the extracellular domain. BigDyn is detected only in UV-exposed samples containing AzF in the acidic pocket, but absent in control samples not exposed to UV, those carrying AzF in the lower extracellular domain (right insets in (A)) or WT. See also Figure S7.

## Discussion

### Mechanism of action of BigDyn on ASIC1a

Despite the potential pathophysiological relevance of the ASIC-BigDyn interaction (10), the determinants for the high-potency modulation of ASIC steady-state desensitization by BigDyn have remained enigmatic. A defining characteristic of neuropeptide signaling is its slow time scale relative to the fast transmission achieved by small-molecules (1, 2). Indeed, we and others have found that the functional effects of BigDyn on ASIC1a-mediated currents require relatively long preincubation times ((10) and Figure 1). The notion of a slow binding and unbinding process is directly supported by our VCF data, which demonstrate that the BigDyn-induced conformational changes are concentration-dependent and drastically slower than those induced by changes in proton concentration (Figure 1, Figure S2B, Figure 3E). Once bound, BigDyn reduces the proton sensitivity of both pH-dependent activation and steady-state desensitization of ASIC1a. Although the acidic pocket is not likely to be the primary proton sensor of ASICs (37, 38), side chains in and around the acidic pocket have been shown to modulate proton sensitivity (30, 37), and X-ray crystallographic data show substantial contraction of this pocket, together with changes in side chain interactions, in low pH structures (39). A direct binding of BigDyn to this region is therefore expected to alter proton sensing and channel gating in response to protonation. Interestingly, our VCF data suggest that BigDyn binding favors a resting closed state that is distinct from open and desensitized states induced by protons (or PcTx1) (Figure 1F, Figure 3E). Although we cannot rule out some degree of direct quenching of the fluorophore by BigDyn, several lines of evidence indicate that the fluorescence change is indeed primarily caused by conformational changes: I) high concentrations of DynA/B only have neglectable effects on the fluorescent signal, II) the half-maximal BigDyn concentrations to elicit functional and fluorescence effects are in a similar concentration range, and III) the BigDyn-insensitive Lys105Cys/Glu238Gln/Asp345Asn triple mutant is only marginally quenched by BigDyn at pH 7.0.

We therefore argue that the BigDyn-induced conformation increases the energetic barrier to populate both open and desensitized states, consistent with the right-shift of both parameters in the presence of BigDyn (Figure 1). In the case of steady-state desensitization, this would result in ASIC1a remaining responsive to pH drops even under proton concentrations that would normally desensitize the channel, thus explaining the ability of BigDyn to noticeably increase ASIC1a dependent neuronal death during acidosis (10).

As activation and desensitization both involve the collapse of the acidic pocket, bringing thumb and finger domain aspartate side chains from ∼8 Å to within <3 Å of each other (39), we speculate that BigDyn binding hinders the proton-induced transitions from resting closed states to active and/or desensitized states.

### Defining site of the ASIC1a-BigDyn interaction

Consistent with the hypothesis that the unusually high density of positive charge of BigDyn is crucial for its functional effects, we find that charge-neutralizing mutations of basic amino acids reduce the ability of BigDyn to rescue ASIC1a proton-gated currents after exposure to steady-state desensitization-inducing conditions. This effect was particularly pronounced for mutations in a cluster of three Arg (Arg6, Arg7 and Arg9) in the N-terminal part of the peptide, while no effect was observed when mutating the only non-charged side chain between amino acids 6 and 9 (Ile8). This is in agreement with the finding that DynA (representing the N-terminal part of BigDyn), but not DynB (representing the C-terminal part of BigDyn) can have modulatory effects on ASICs (10, 40) (Figure S1C). However, it is worth noting that functional effects were also reduced when mutating Lys11, Arg19, Arg26 and Tyr1, albeit to a lesser extent (Figure 2 and S3A). These data suggest that the interaction of BigDyn with ASIC1a is not solely driven by charge at Arg6, Arg7 and Arg9, but also mediated by other, mostly basic amino acids throughout the peptide sequence. This notion is further supported by data from our N-terminal truncation screen and complemented by the finding that AzF incorporated at positions along the entire length of the cleft formed by the acidic pocket enabled to covalently crosslink to BigDyn. Together, this argues for an extended interaction surface on ASIC1a, likely extending from the peripheral thumb domain around the α5 helix and into the acidic pocket. While the location of the BigDyn binding site is distinct from that suggested for RFamide neuropeptides on ASICs (41, 42), it does overlap with the binding site for PcTx1 (10, 21, 22, 31). This is consistent with the finding that RFamides do not functionally compete with BigDyn or PcTx1 (10, 43). Interestingly, both PcTx1 and BigDyn bind to the acidic pocket through extensive charge-charge interactions, and additional contributions are made by aromatic amino acids (i.e. Trp7 and Trp24 in PcTx1 (31) and Tyr1 in BigDyn). However, the binding mode is likely to differ substantially between the two peptides, as PcTx1 adopts a rigid fold, while BigDyn is likely unstructured in aqueous solution (44, 45). Indeed, the two peptides have opposite modulatory effects on ASIC1a: while PcTx1 interacts with and stabilizes the desensitized state (27), our data suggest that BigDyn stabilizes a closed/resting state of ASIC1a (see above).

Future studies using computational docking, ligand binding studies or a combination of crosslinking and mass spectrometry could further help to provide details on the precise binding mode and orientation of BigDyn at the acidic pocket.

### Possible biological implications

Intriguingly, both ASIC1a and dynorphins have been shown to exhibit overlapping expression patterns (e.g. amygdala, hippocampus) and to contribute to a similar array of both physiological (learning, memory) and pathophysiological (pain, nociception, addiction) processes (4, 5, 11, 12, 46–52). Among the dynorphins tested here, BigDyn is the most potent at ASIC1a and has also been shown to greatly enhance acidosis-induced cell death in cortical neurons (10). The potential relevance of this finding is underscored by the fact that under pathophysiological conditions, the BigDyn levels will be sufficient to modulate ASIC activity *in vivo* (up to low micromolar range) (4, 5, 8, 10, 53, 54). By contrast, the modulatory effects under physiological BigDyn concentrations (1-10 nM) would likely be small, although others have reported a higher sensitivity of ASIC1a towards BigDyn (10). Together, this highlights the potential of the ASIC-BigDyn interaction site as a drug target. Given that BigDyn and PcTx1 share a common binding site, this notion is supported by the finding that PcTx1 or PcTx1-like peptides show promise in limiting neuronal death in stroke models (14, 15). While the ASIC-BigDyn interaction *per se* increases neuronal death, a therapeutic option might in the future emerge from building upon the scaffold of modified peptides with reduced activity (similar to some presented in this study) that could, if refined, work as silent modulators to prevent the ASIC–BigDyn interaction. Recent work on PcTx1 has shown that peptides targeting the ASIC acidic pocket can display significant functional plasticity. For example, altered pH or mutations within the toxin (Arg27Ala or Phe30Ala) can effectively convert PcTx1 into a potentiator (31, 55). Similarly, and depending on cellular context and presence of co-factors, DynA can act as both an inhibitor and potentiator of glutamate receptors (4). It will thus be intriguing to see if this functional dichotomy is also possible to achieve for BigDyn in order to unlock its potential as an ASIC-targeting therapeutic lead.

## Methods

Mouse ASIC1a (mASIC1a) was expressed in *Xenopus laevis* oocytes and proton induced currents were measured with two-electrode voltage clamp electrophysiology (TEVC). Synthetic dynorphin peptides and analogs were tested for their ability to decrease the pH sensitivity and rescue currents of mASIC1a from proton induced steady-state desensitization (Fig. 1, 2). Proton and peptide induced conformational rearrangement in mASIC1a were monitored by voltage-clamp fluorometry (VCF) (Fig. 1E-G). Site-directed mutagenesis was employed to introduce charge-neutralizing amino acid substitutions of negatively charged residues of the acidic pocket and the sensitivity of mutant mASIC1a towards BigDyn was assessed by TEVC (Fig. 3). UV crosslinking of BigDyn to human ASIC1a (hASIC1a) was achieved in HEK 293T cells in which endogenous hASIC1a was removed by CRISPR/Cas9 using guide RNA (56). ASIC1a-free HEK 293T cells were transfected with DNA encoding hASIC1a or hASIC1a TAG variants and AzF was introduced through the non-sense suppression method (35). 3 µM BigDyn were added to AzF-hASIC1a expressing HEK 293T cells and the cells were subsequently exposed to UV-light for 15 min in order to induce crosslinking. UV-treated cells were lysed and the UV-induced hASIC1a-BigDyn complex formation was visualized by western blotting (Fig. 4). See SI Appendix, Supplementary Text for detailed description of materials and methods.

## Acknowledgements

We acknowledge the Lundbeck Foundation (R139-2012-12390 to SAP and R218-2016-1490 to NB), the Boehringer Ingelheim Fond (to NB), the Danish National Research Foundation (DNRF107 to WT), the European Union’s Horizon 2020 research and innovation program under the Marie Sklodowska-Curie grant agreement No 834274 (to SAH) and the University of Copenhagen for financial support.

## Author contributions

CBB, NB, SAH, EPB, LMHK, TL, KS, JA and SAP designed the research; CBB, NB, SAH, YB, DW, IG, CL, TW and LMHK performed the research and analyzed the data; JA and SAP supervised the project; CBB, NB, SAH and SAP wrote the manuscript with input from all authors.

## Competing interests

The authors declare to have no competing interests.

## Supplementary Information

### Supplementary Text

#### Materials and Methods

##### Molecular biology

The complementary DNA (cDNA) encoding mouse ASIC1a (mASIC1a) was used as previously described (1), while the human ASIC1a (hASIC1a) cDNA was kindly provided by Dr Stephan Kellenberger. Plasmids containing AzF-RS and tRNA were gifts from Dr Thomas P. Sakmar (2). The dominant negative eukaryotic release factor (DN-eRF) was a gift from Dr William Zagotta (3).

Site-directed mutagenesis was performed using PfuUltraII Fusion polymerase (Agilent) and custom DNA mutagenesis primers (Eurofins Genomics). All sequences were confirmed by sequencing of the full coding frame (Eurofins Genomics). For hASIC1a constructs, a C-terminal 1D4-tag was added for protein purification and Western blot detection and two silent mutations were inserted at V10 and L30 to reduce the risk of potential reinitiation (4). cDNAs were linearized with EcoRI (mASIC1a) or HindIII-HF (hASIC1a, both New England Biolabs) and capped cRNA was transcribed with the Ambion mMESSAGE mMACHINE SP6 (mASIC1a) kit (Thermo Fisher Scientific).

##### Electrophysiological recordings

Oocytes were surgically removed from adult female *Xenopus laevis* and prepared as previously described (1). Oocytes were injected with 0.1 to 10 ng cRNA of mouse ASIC1a (mASIC1a) or hASIC1a mRNA into the cytosol (volumes between 9 and 50 nL). Typically, higher amounts of mutant mASIC1a constructs were injected compared WT mASIC1a. 1-4 days after injection, oocytes were transferred to a recording chamber (5), continuously perfused (2.5 mL/min) with bath solution containing (in mM): 96 NaCl, 2 KCl, 1.8 BaCl_2_, 2 MgCl_2_, and 5 HEPES, pH adjusted by NaOH or HCl. Solutions exchange was achieved using a gravity-driven 8-line automated perfusion system operated by a ValveBank module (AutoMate Scientific). Current recordings were performed in presence of 0.05% bovine serum albumin (≥ 98% essentially fatty acid-free, Sigma Aldrich).

Currents were recorded using microelectrodes (borosilicate capillaries 1.2 mm OD, 0.94 mm ID, Harvard Apparatus), backfilled with 3 M KCl (0.5-2 MΩ) and an OC-725C amplifier (Warner Instruments). The current signal was acquired at 500 Hz, filtered by a 50-60 Hz noise eliminator (Hum Bug, Quest Scientific) and digitized using an Axon^TM^ Digidata^®^ 1550 Data Acquisition System and the pClamp (10.5.1.0) software (Molecular Devices). The current signal was further digitally filtered at 2.8 Hz using an 8-pole Bessel low-pass filter prior to analysis. Displayed current traces have been subjected to additional 50× data reduction.

Synthetic PcTx1 was obtained from Alomone Labs (>95 % purity). PcTx1 and synthetized dynorphin peptides and analogs stock solutions were prepared in MilliQ (18.2 MΩ resistivity). Peptide stock solutions were stored at −20 °C. Prior to recording, the peptides were diluted to the desired concentration in recording solution.

Concentration-response relationships of WT mASIC1a activation were determined by 20 s applications of solutions with decreasing pH-values. Between each 20 s application, the oocytes were left to recover for 2 min in pH 7.4 solution (unless stated otherwise). The currents elicited by the 20 s application of acidic pH were normalized to the largest current size for each of individual oocyte tested. Steady-state desensitization concentration-response relationships were determined by exposing ASIC1a expressing oocytes to a 20 s application of pH 5.6 (unless stated otherwise), while decreasing the resting pH in between applications of pH 5.6. The resting period between pH 5.6 applications was 2 min. In order to ensure that the observed current desensitization was not due to general current run-down, a final application of pH 5.6 was performed after a 2 min resting period at a resting pH that resulting in saturating current responses. Traces were used for further analysis only if the final pH 5.6 application resulted in currents that were ≥ 80 % of the same resting pH prior to the steady-state desensitization protocol.

The ability of dynorphin peptides to rescue ASIC1a proton-gated currents after exposure to steady-state desensitization-inducing conditions was examined at a resting pH of 7.6 (unless stated otherwise). Channel activation was achieved by application of pH 5.6 (except for Asp237Asn/Asp345Asn and Glu373Gln/Glu411Gln constructs which required pH 4.0 for full activation) for 20 s. The oocytes were left to recover for 1 min in pH 7.6 solution. Hereafter, the oocyte was perfused for 2 min with steady-state desensitization-inducing pH (determined specifically for each individual ASIC1a construct/mutant) prior to a 20 s application of pH 5.6. Dynorphin peptides were applied during the 2 min of steady-state desensitization conditioning. The channels were checked for ability to recover (cutoff at ≥80 % recovery) after 3 min of pH 7.6 solution by an additional 20 s application pH 5.6. The effect of the dynorphin peptides was determined by normalizing the current post-peptide application to the mean of the two currents elicited without steady-state desensitization conditioning pH.

##### Voltage-clamp fluorometry

For voltage-clamp fluorometry (VCF), 5-20 ng of mASIC1a Lys105Cys mRNA were injected into oocytes which were then incubated for 2-7 days. On the day of the recording, oocytes were labeled by incubating them for about 30 min with 10 μM AlexaFlour 488 C5-maleimide (Thermo Fisher Scientific) in OR2 solution at room temperature, subsequently washed twice with OR2, and stored in the dark until further use. The oocytes were placed in the custom-built recording chamber above a water immersion objective of an inverse microscope (IX73 with LUMPlanFL N 40x, Olympus), with the animal pole facing the excitation source. An Olympus TH4-200 halogen lamp or a 470 nm CoolLed pE-100 excitation light source with a standard GFP filter-set (Olympus) was used to excite the fluorophore while the emission was detected using a P25PC-19 photomultiplier tube (Sens-Tech) and photon-to-voltage converter (IonOptix), via the microscope side port. Current and fluorescence signals were acquired, filtered, digitized and digitally filtered as described for the other electrophysiology recordings.

In all VCF experiments, except the ones described In Figure S2A, 0.05% BSA was added to all solutions (composition as described above). Channels were activated for 20 s using pH 5.5 buffer, washed for 1 min with pH 7.6 buffer, exposed for up to 2 min to 0.3 µM PcTx, 1–10 µM BigDyn, DynA or DynB, or pH 9.0, followed by a 2 min washout at pH 7.6 and a subsequent 20 s exposure to pH 5.5. For the pH response curve in SI Figure S2A, oocytes were exposed for 20 s to pH 6.0, washed for 1 min at pH 7.6, exposed for 1-2 min (until fluorescent signal reached a plateau) at pH 7.2, followed by direct exposure to pH 6.0. After a 1 min washout at 7.6 this protocol was consecutively repeated using decreasing preconditioning pH (7.1, 7.0, 6.8, 6.6), in the end the response to pH 6.4 was measured. All signals were reported relative to deflections at pH 6.0 for the pH-response curve in SI Figure S2, relative to the 10 µM BigDyn deflection in Figure S2B-C, or relative to deflections at pH 5.5 for all other experiments. Fluorescent signal intensity was reported as percentage change relative to the baseline fluorescence level and the fluorescent baseline was adjusted for all fluorescence traces.

##### Cell culture and transfection

HEK 293T cells (ATCC^®^) were grown in monolayer in T75 and T175 flasks (Orange Scientific) in DMEM (Gibco) supplemented with 10 % FBS (Thermo Fisher Scientific) and 1 % penicillin-streptomycin (Thermo Fisher Scientific) and incubated at 37 °C in a humidified 5 % CO_2_ atmosphere. Endogenous hASIC1a was removed by CRISPR/Cas9 using the guide RNA published in (6). For crosslinking studies, cells were seeded into 15 cm dishes (VWR) at a density of 5-7 mio. cells and transfected the next day with PEI (Polysciences) and 16:4:4:8 µg DNA encoding hASIC1a TAG variants, AzF-RS, tRNA and DN-eRF, respectively. For the WT control, 2 mio. cells were seeded into a 10 cm dish (VWR) and transfected with 8 µg hASIC1a WT. Six hours after transfection, cell medium was replaced with DMEM containing 0.5 mM AzF (4-Azido-L-phenylalanine, Chem Impex) and crosslinking studies were performed 48 hours after transfection.

##### Crosslinking studies, protein purification, western blotting

Cells were washed with PBS and dislodged from the dishes using cell scrapers (Orange Scientific). After centrifugation at 1000 rpm for 5 min, cell pellets were resuspended in 1 ml PBS containing 3 µM BigDyn (pH 7.4) and transferred into 12 well plates (Orange Scientific). Cells were crosslinked under a Maxima ML-3500 S UV-A light source (Spectronics corporation, 365 nm) for 15 min on ice. Control samples without UV exposure were kept at 4 °C. After crosslinking, cells were centrifuged at 1000 rpm for 5 min and resuspended in 1 ml solubilisation buffer (50 mM Tris-HCl, 145 mM NaCl, 5 mM EDTA, 2 mM DDM, pH 7.5) supplemented with cOmplete™ EDTA-free protease inhibitor cocktail (Sigma Aldrich). Cells were lysed at 4 °C for 2 h and centrifuged at 18000 g and 4 °C for 30 min. In parallel, 40 µl Dynabeads Protein G (Thermo Fisher Scientific) were washed with 200 µl PBS/0.2 mM DDM and incubated with 4 µg RHO 1D4 antibody (University of British Columbia) in 50 µl PBS/0.2 mM DDM for 30 min on a ferris wheel (VWR). After washing the beads with 200 µl PBS/0.2 mM DDM, the cell lysate supernatant was incubated with the beads on the ferris wheel at 4 °C for 90 min. Beads were washed with 200 µl PBS three times to remove nonspecifically bound proteins and incubated in 25 µl elution buffer (2:1 mixture between 50 mM glycine, pH 2.8 and 62.5 mM Tris-HCl, 2.5 % SDS, 10 % Glycerol, pH 6.8) supplemented with 80 mM DTT at 70 °C for 10 min. Protein samples (12 µl) were mixed with 3 µl 5 M DTT and 5 µl 4x NuPAGE™ LDS sample buffer (Thermo Fisher Scientific) and incubated at 95 °C for 20 min before SDS-PAGE using 3–8 % Tris-Acetate protein gels (Thermo Fisher Scientific). After transfer onto PVDF membranes (iBlot 2 Dry Blotting System, Thermo Fisher Scientific) and blocking in TBST/3% non-fat dry milk for 1 hour, hASIC1a was detected using RHO 1D4 antibody (1 µg/µl, University of British Columbia) and 1:5000 goat anti-mouse IgG HRP-conjugate (Thermo Fisher Scientific). BigDyn was detected using 1:1000 rabbit anti-DynA antibody (abcam) and 1:5000 goat anti-rabbit IgG HRP-conjugate (Promega).

##### Solid-phase peptide synthesis

Unless otherwise stated, all amino acids and reagents were purchased from either Iris Biotech, Gyros Protein Technologies or Sigma Aldrich. All solvents were purchased from commercial sources and used without further purification. The molecular mass of the peptides spectra was obtained by electron spray ionization (ESI) liquid chromatography mass spectrometry (LC-MS) coupled to an Agilent 6410 Triple Quadrupole Mass with a reverse phase C18 column (Zorbax Eclipse XBDC18, 4.6 × 50 mm) using a binary buffer system consisting of H_2_O:CH_3_CN:formic acid (A, ratio 95:5:0.1; B, ratio 5:95:0.1) at 0.75 mL/min. Peptide purity was determined by UV absorbance at 214 nm on an analytical reverse phase ultraperformance liquid chromatography (RP-UPLC) (Waters) system with a C18 column (Acquity UPLC BEH C18, 1.7 μm 2.1 × 50 mm) using a binary buffer system consisting of H_2_O:CH_3_CN:trifluoroacetic acid (TFA) (A, ratio 95:5:0.1; B, ratio 5:95:0.1) at 0.45 mL/min.

Peptides were synthesized by Fmoc solid-phase peptide synthesis (SPPS) at 0.04-0.10 mmol scale. For SPPS, preloaded 4-benzyloxybenzyl alcohol (Wang) (Iris Biotech), TentaGel^®^ R PHB or preloaded Fmoc-Thr(OtBu) TentaGel^®^ R PHB (Rapp Polymere) resins (100-200 mesh) were used. Loading of hydroxy-functionalized TentaGel^®^ R PHB resin was carried out by the MSNT/MeIm (1-(2-mesitylenesulfonyl)-3-nitro-1H-1,2,4-triazole/1-methylimidazole) method (7). Loading of resin was checked spectrophotometrically, quantifying the amount of Fmoc released upon cleavage as previously described (8). Standard coupling reactions were carried out at room temperature (RT) under agitation using a mixture of Fmoc-protected amino acid derivative (4.0 equiv. relative to resin loading), N,N,N**′**,N**′**-tetramethyl-O-(1H-benzotriazol-1-yl)uranium hexafluorophosphate (HBTU, 4.0 equiv.) and N,N-diisopropylethylamine (DIPEA) (8.0 equiv.) in dimethylformamide (DMF). Coupling reactions were monitored using a Kaiser test kit (Sigma Aldrich). Fmoc deprotections were carried out by treatment with 20 % piperidine in DMF (2 × 2 min). After coupling or Fmoc deprotection steps, the resin was extensively washed with DMF.

Automated peptide synthesis was carried out using either a Prelude^®^ X peptide synthesizer equipped with induction heating (Gyros Protein Technologies) or using a Syro Wave^TM^ Parallel Peptide Synthesizer (Biotage). For Prelude X synthesis all reagents were prepared in DMF: Fmoc-protected amino acid (0.2 M), O-(1H-6-chlorobenzotriazole-1-yl)-1,1,3,3-tetramethyluronium hexafluorophosphate (HCTU) (0.5 M) DIPEA (1 M) and Fmoc deprotection solution (20% piperidine, v/v). Sequence elongation was achieved by following protocol: Fmoc deprotection (2 × 2 min, 20 % piperidine (v/v), rt,) and coupling (2 × 5 min, 75 °C, for Cys couplings 2 × 5 min, 50 °C, and for Arg couplings 3 × 5 min, 50 °C). Amino acids were double coupled using amino acid:HCTU:DIPEA (ratio 1:1.25:2.5) in 5-fold excess over the resin loading to ensure efficient peptide elongation. Final Fmoc deprotection after peptide elongation (2 × 5 min) was followed by dichloromethane (DCM) wash.

For Syro Wave^TM^ synthesis Fmoc-amino acids (0.5 M), HBTU (0.48 M) and piperidine (40 % v/v) were prepared in DMF. DIPEA (2 M) was prepared in N-methyl-2-pyrrolidone (NMP). Sequence elongation was achieved using following protocol: Fmoc deprotection (1 × 3 min treatment with 40 % piperidine in DMF followed by 1 × 12 min treatment in 20 % piperidine) and coupling (1 x 40 min, RT, for Arg and Cys couplings 2 x 40 min, RT). Amino acids were coupled using amino acid:HBTU:DIPEA (ratio 1:1:2) in 5-fold excess over the resin loading. After deprotection, the resin was washed with DMF (× 2), DCM (× 1), and DMF (× 2) for 1 min each. The final Fmoc-deprotection was achieved by treatment with 40 % piperidine in DMF (3 min), followed by treatments with 20 % piperidine in DMF (2 × 8 min).

Global deprotection and cleavage was conducted in a mixture of TFA:triisopropylsilane (TIPS):H_2_O (ratio 95:2.5:2.5, v:v:v) for 2 h at room temperature. Subsequently, the cleaving mixture was concentrated and the crude peptide was precipitated with ice-cold diethyl ether. The precipitate was centrifuged at 3500 x g, 5 min at 4 °C, washed with ice-cold diethyl ether and centrifuged at 3500 x g, 5 min at 4 °C, after which it was solubilized in H_2_O:CH_3_CN:TFA (ratio 50:50:0.1, v:v:v) and lyophilized.

Peptides were purified using either a preparative reversed-phase high-performance liquid chromatography (RP-HPLC) system (Waters) with a RP C18 column (Zorbax, 300 SB-C18, 7 µm, 21.2 × 250 mm) or an Agilent 1260 LC system equipped with a diode array ultraviolet detector and an evaporative light-scattering detector (ELSD) using a RP-C8 column (Phenomenex Luna, 5 μm, 100 Å, 21.2 × 250 mm). A linear gradient with a binary buffer system of eluent A (H_2_O:CH_3_CN:TFA, ratio 95:5:0.1, v:v:v) and eluent B (CH_3_CN:TFA, 99.9:0.1, v:v) was applied, with a flow rate of 20 mL/min. Collected fractions were characterized by LC-MS. Pure fractions were pooled together and lyophilized, which yielded a white, fluffy material. The purity of final peptides was determined by RP UPLC monitoring the absorbance at 214 nm and the results are listed in Table S8.

##### Data analysis

Data analyses were performed in Prism (8.0) (GraphPad Software). For proton concentration-response data, peak current amplitudes were plotted against pH and fitted with the Hill equation for each recording. These were averaged to give the reported means ± 95CI in the main text. For display in figures, a single fit to the average normalized responses (± 95CI) is shown. All bar diagrams and summarized data points are presented as mean ± 95CI unless stated otherwise and number of replicates (n) represents individual experimental oocytes. Results were obtained from at least two batches of oocytes unless indicated differently. An unpaired t-test was used to compare two groups. Multiple comparisons were made with one-way analysis of variance with Dunnett’s comparison to a control value (e.g. comparing with WT) or with Tukey’s test for multiple comparisons. A significance level of p < 0.05 was applied for all analyses. All graphs and illustrations were made in Prism (8.0) (GraphPad Software) and Illustrator CC (Adobe).

## Supplementary Figures

**Figure S1:**
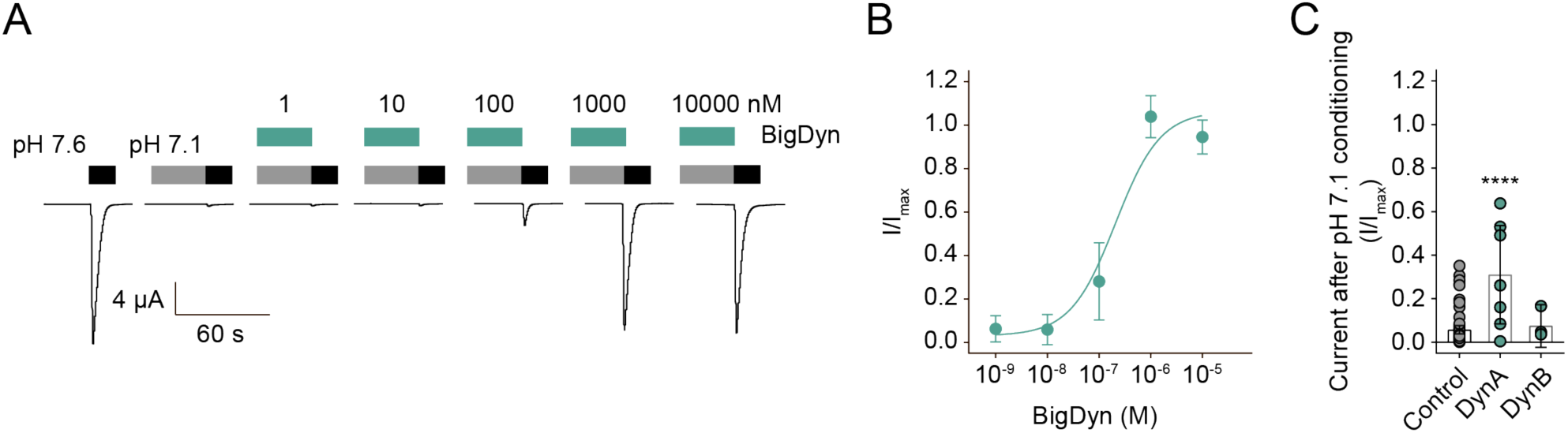
Modulation of ASIC1a by BigDyn, DynA and DynB. (A/B) Representative current traces (A) and normalized data (B) obtained by pH 5.6 application (black bar in (A)) at ASIC1a WT-expressing *Xenopus laevis* oocytes after preincubation in pH 7.1 (grey bar) with or without different concentrations of BigDyn (green bars). EC_50_ of BigDyn is 210 nM (95CI: 163, 259 nM); in (A) pH 5.6 current response after pH 7.6 incubation is shown as control; data shown as mean ± 95CI; n = 9. (C) Averaged data obtained by pH 5.6 application at mASIC1a WT after preincubation in pH 7.1 (grey bar) with or without 1 µM of the indicated peptide (green bar); each dot represents an individual oocyte and bars represent mean + 95CI. Asterisk in (C) indicates significant difference to control condition (p < 0.0001) n = 4-68.

**Figure S2:**
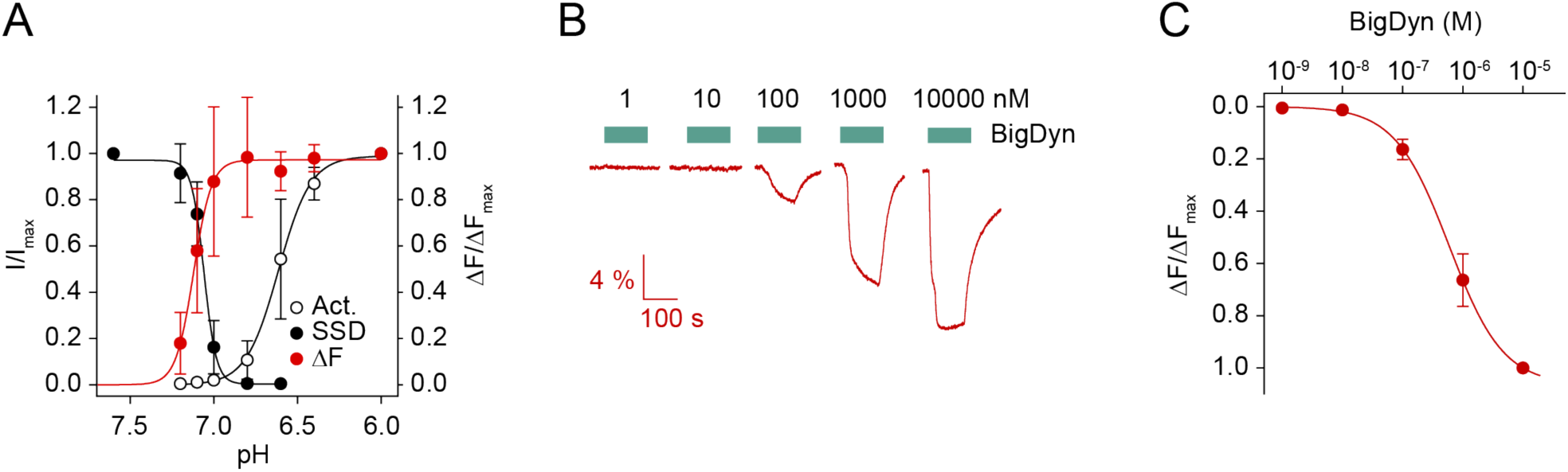
Proton dependence of current and fluorescence signals generated by the Alexa 488-labelled Lys105Cys mutant. (A) Normalized current activation (Act.) and steady-state desensitization (SSD, black dots, left y-axis) and fluorescence (red dots, right Y-axis) signals are plotted as a function of pH. For ΔF, the fitting was constrained using min. = 0. Data shown as mean ± 95CI; n = 5. (B) Representative fluorescence traces obtained by application of increasing BigDyn concentrations. (C) Normalized change in fluorescence plotted as a function of BigDyn concentration as seen in (B). The fitting was constrained using max. = 0. All data is obtained from *Xenopus laevis* oocytes. Data shown as mean ± 95CI; n = 9.

**Figure S3:**
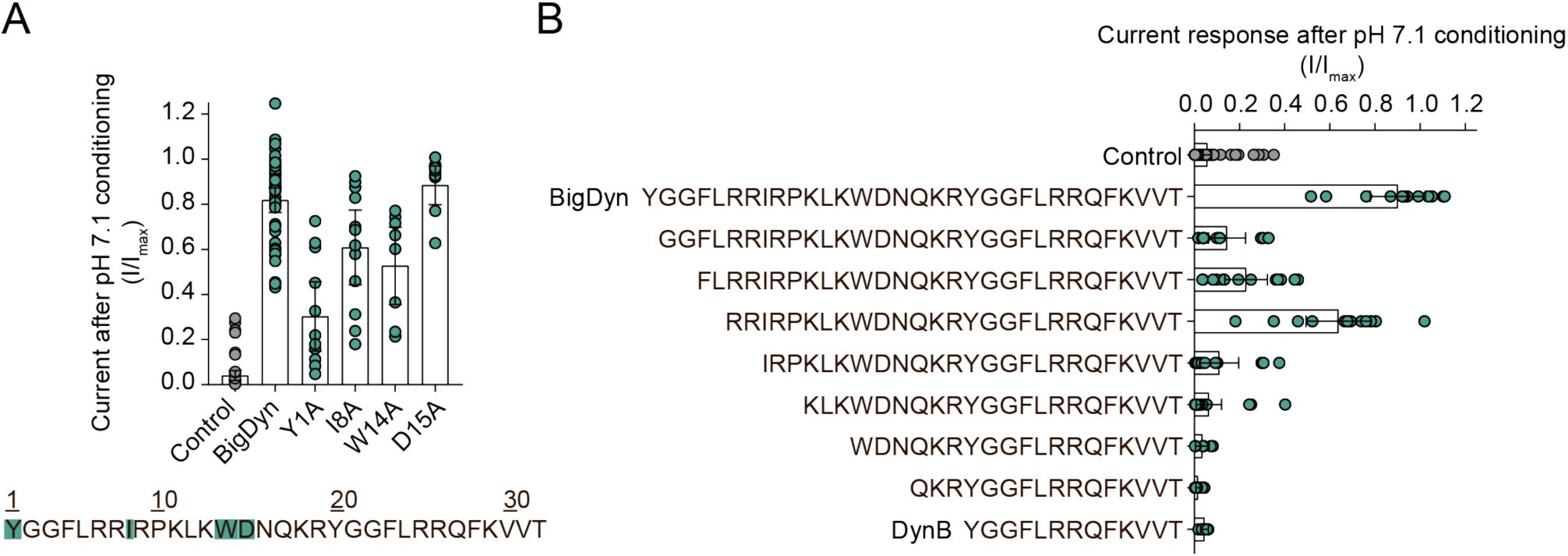
Modulation of ASIC1a by BigDyn derivatives. (A) Averaged data obtained by pH 5.6 application at mASIC1a WT-expressing *Xenopus laevis* oocytes after preincubation in pH 7.1 in the presence of 1 µM of the indicted peptide. Each dot represents an individual oocyte and bars represent mean ± 95CI, n = 10-48. The BigDyn sequence with side chains mutated to alanine highlighted in green is shown below the graph. (B) Averaged data obtained for different N-terminal truncation variants of BigDyn obtained by plotting pH 5.6-induced current amplitude after preincubation on pH 7.1 at ASIC1a WT in the presence of 1 µM of the indicated peptide. Each dot represents an individual oocyte and bars represent mean ± 95CI; n = 6-68.

**Figure S4:**
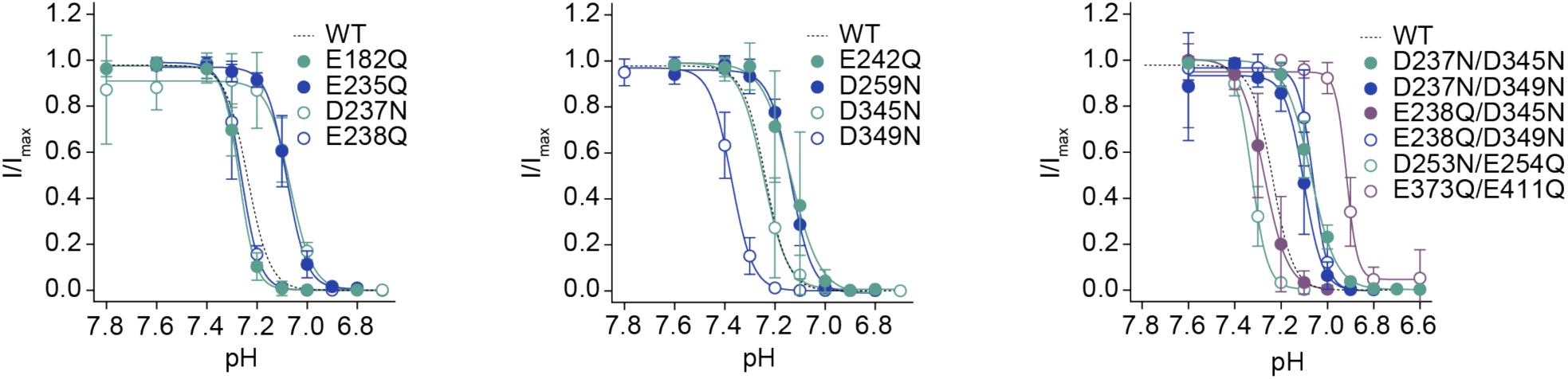
Steady-state desensitization of single and double charge-neutralizing ASIC1a mutants. Normalized steady-state desensitization plotted as a function of pH. Data shown as mean ± 95CI; n = 3-64. See Table S6 for further details.

**Figure S5:**
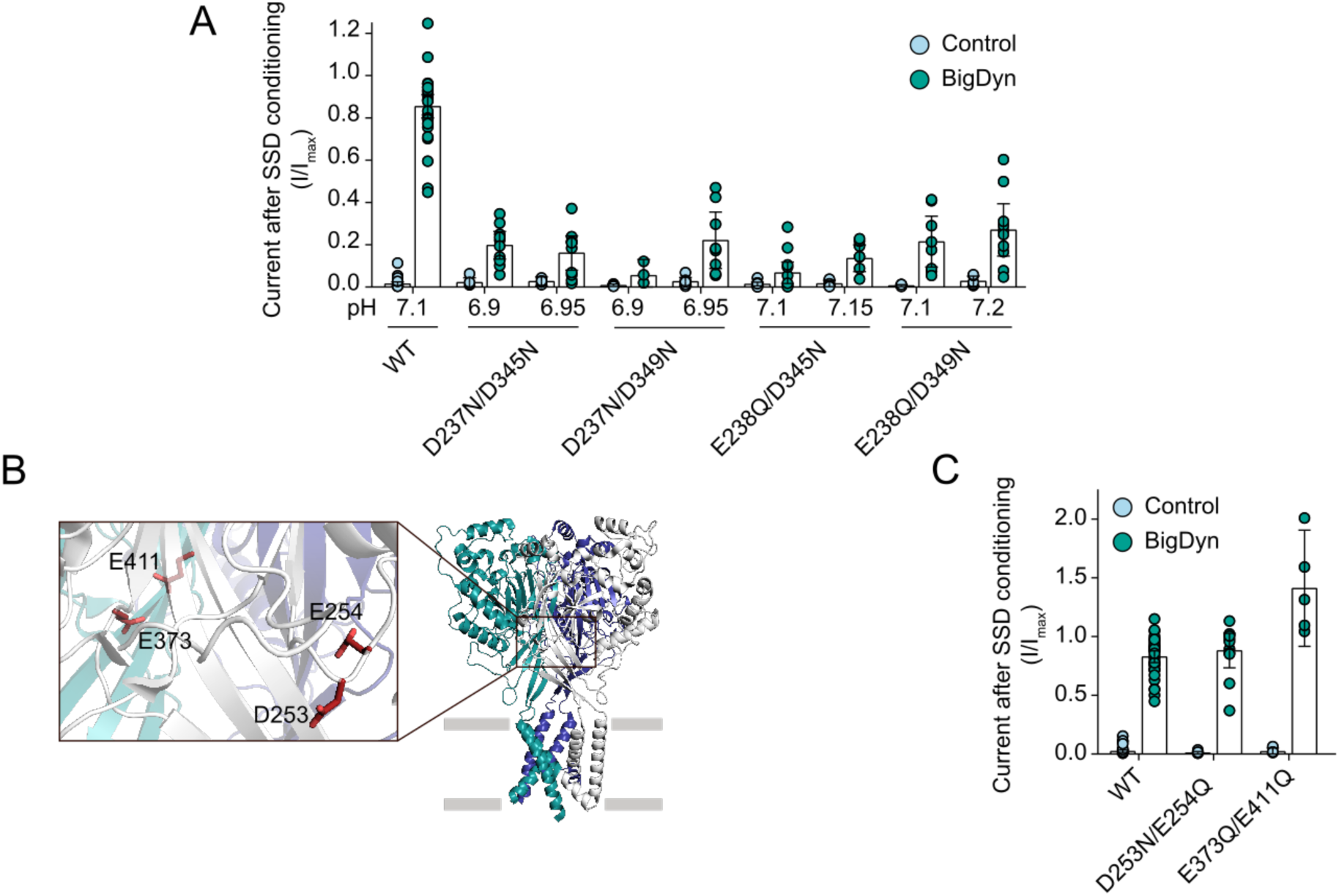
Effects of BigDyn on double charge-neutralizing mutations. (A) Averaged data obtained by pH 5.6 application after preincubation in different steady-state desensitization (SSD)-inducing pH conditions with or without (Control) 1 µM BigDyn at *Xenopus laevis* oocytes expressing the indicated double mutant ASIC1a construct; n = 4-33. (B) Structure of cASIC1a (PDB: 4NTW) with individual subunits color coded and inset showing the location of acidic side chains away from the acidic pocket that have been mutated to glutamine or asparagine. (C) Averaged data obtained by pH 5.6 (WT and Asp253Asn/Glu254Gln) or 4.0 (Glu373Gln/Glu411Gln) application after preincubation in steady-state desensitization-inducing pH with or without (Control) 1 µM BigDyn at *Xenopus laevis* oocytes expressing the indicated double mutant ASIC1a construct; n = 5-45. Each dot in (A), (C) represents an individual oocyte and error bars represent mean ± 95CI.

**Figure S6:**
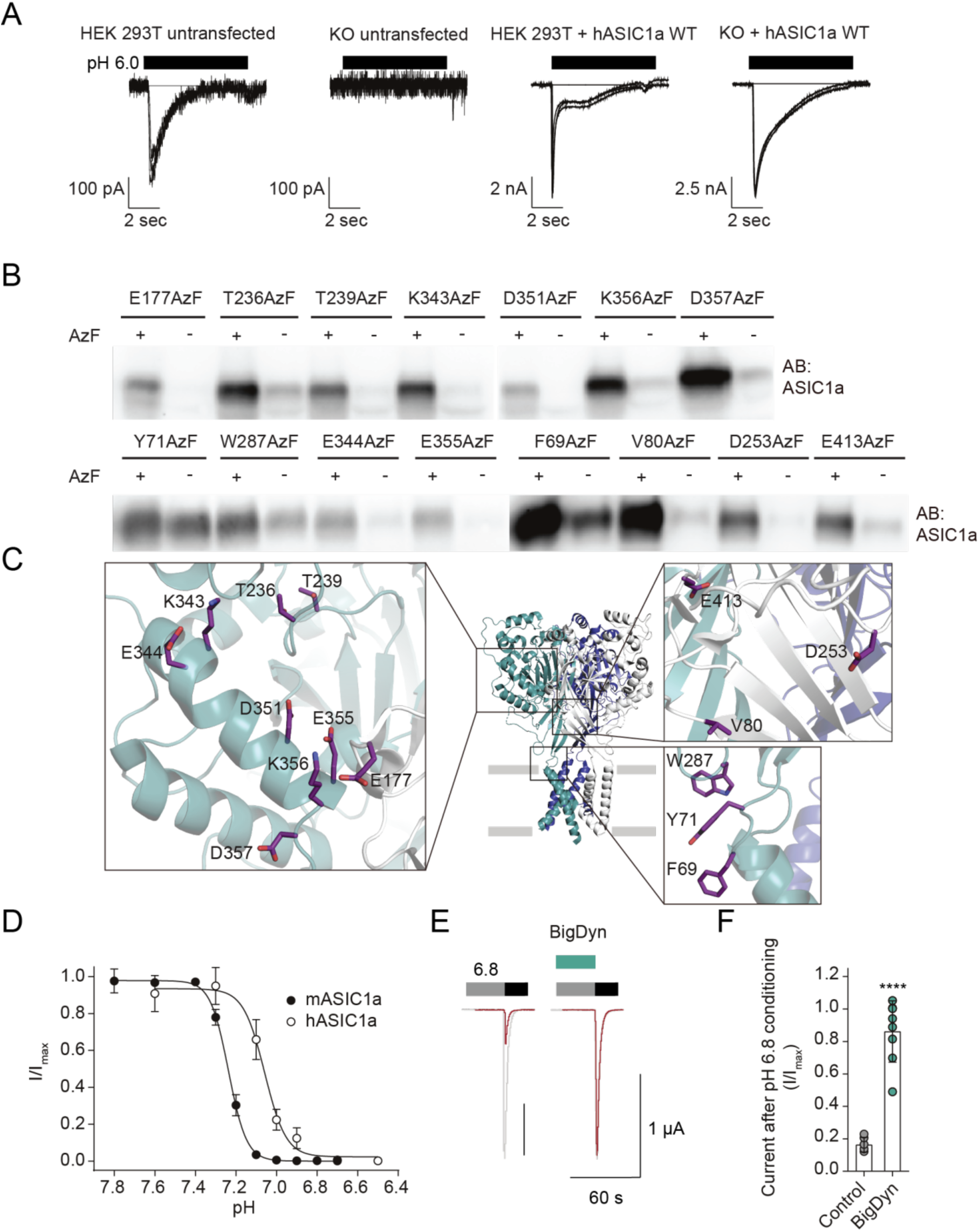
Incorporation of AzF and BigDyn modulation of hASIC1a. (A) Proton-gated currents in untransfected HEK 293T cells are abolished by CRISPR/Cas9 KO of endogenous hASIC1a and rescued through transfection of hASIC1a WT. pH 6.0 induced currents were measured on a SyncroPatch 384PE (Nanion Technologies, whole-cell patch-clamp, n = 5-12 cells). Note that the differences in current shape are likely due to the ligand application on the SyncroPatch 384PE. (B) Western blot analysis of the indicated stop-codon containing mutants grown in the presence or absence of 0.5 mM AzF in HEK 293T cells. After 48 hrs, the resulting full-length protein was purified via a C-terminal 1D4-tag and visualized by western blotting. With the exception of positions 69 and 71, all positions tested here showed only small amounts of full length protein in absence of AzF (compared to those obtained in the presence of AzF), demonstrating efficient incorporation. (C) Structure of cASIC1a (PDB: 4NTW) with individual subunits color coded and insets highlighting side chains replaced by AzF in the acidic pocket (left inset) and lower extracellular domain (right insets). (D) Steady-state desensitization of hASIC1a (open, pH_50_ steady-state desensitization 7.07 (95CI: 7.03, 7.11)) is shifted compared to mASIC1a (filled, pH_50_ steady-state desensitization 7.24 (95CI: 7.23, 7.25, n = 6-8 oocytes)). (E/F) Representative current traces (E) and averaged data (F) comparing the normalized pH 5.6-gated current (black bar) remaining after conditioning hASIC1a at desensitizing pH 6.8 (grey bar) in presence (green bar) and absence of 1 µM BigDyn. Data in F is shown as mean ± 95CI. Asterisk indicates significant difference to control condition (p < 0.0001), n = 6-8 oocytes.

**Figure S7:**
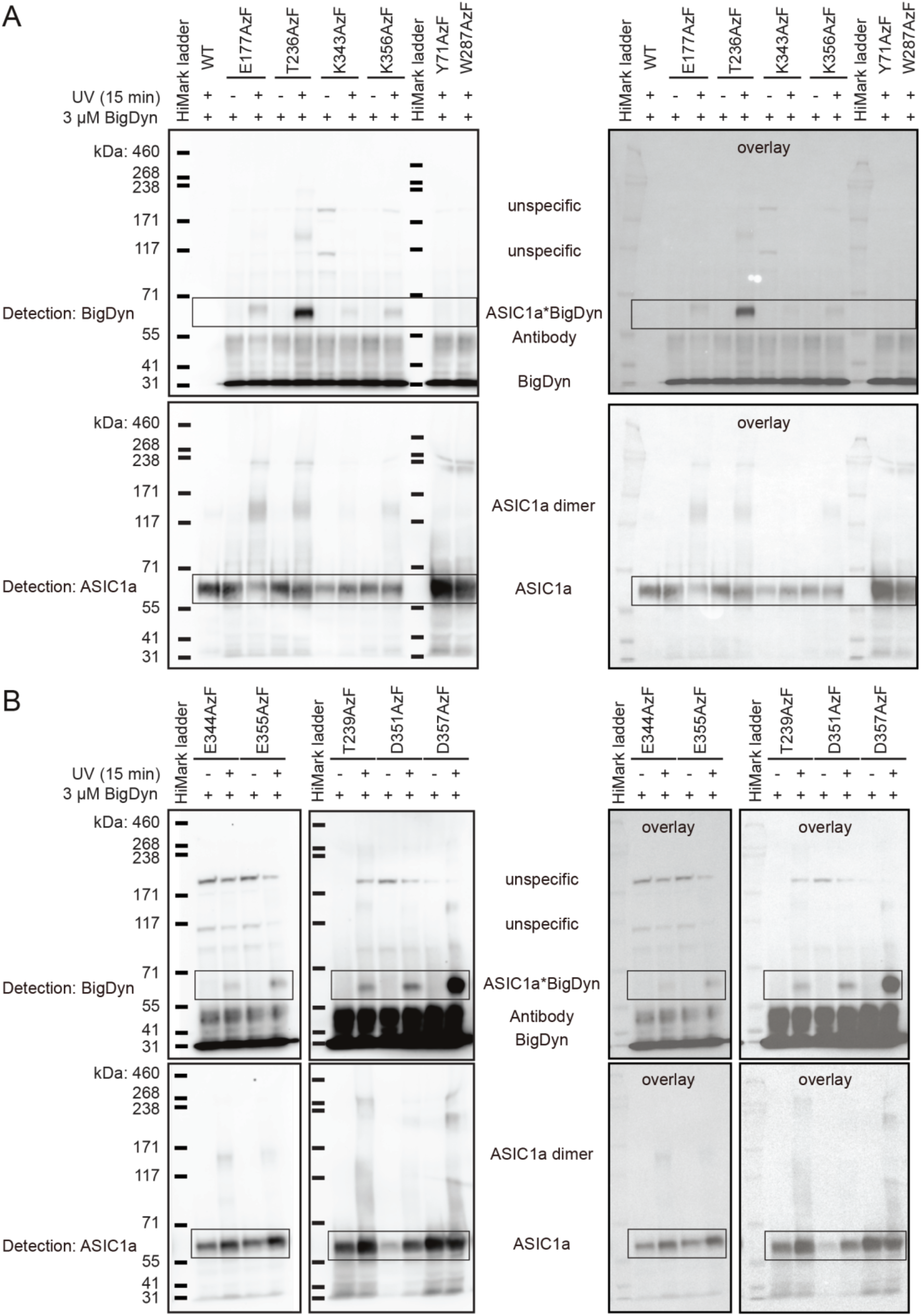
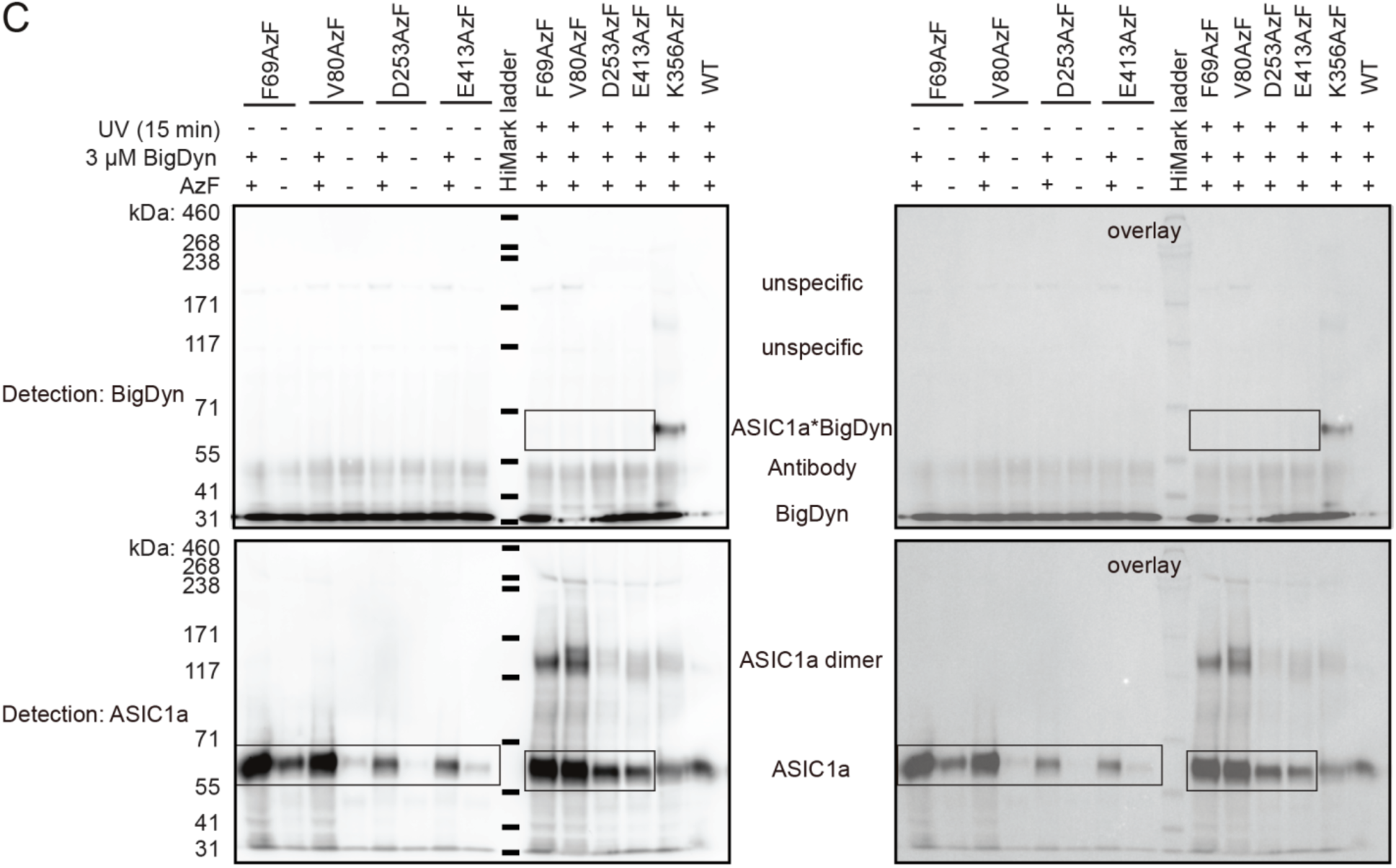
Original Western blots. Left panel contains black bars indicating protein ladders for visibility, right panel shows overlay of original marker (coomassie) and blot (chemiluminescence). Areas cropped for Figure 4 are marked with boxes. (A) Western blot for positions 177, 236, 343, 356, 71 and 287. (B) Western blot for positions 344, 355, 239, 351 and 357. (C) Western blot for positions 69, 80, 253 and 413. Data is representative of 2-3 individual experiments.

**Table S1:**
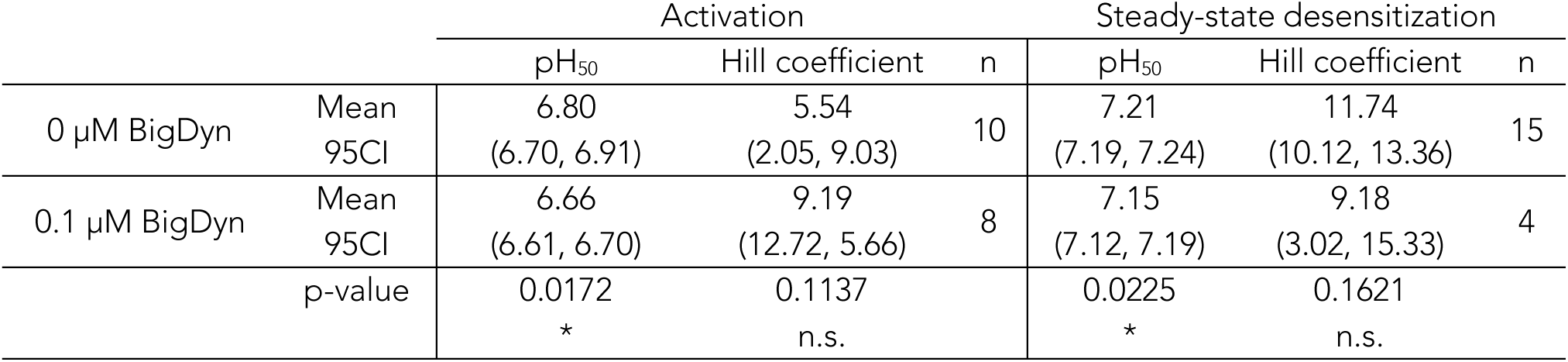
Activation and steady-state desensitization in absence and presence of BigDyn. pH_50_ and Hill coefficients for activation and steady-state desensitization of WT mASIC1a in the absence and presence of 0.1 µM BigDyn. 95CI in parentheses; n = number of experiments; pH_50_-values and Hill coefficients in presence and absence of BigDyn was compared using an unpaired t-test, asterisks indicate statistically significant differences, n.s = not significant.

**Table S2:**
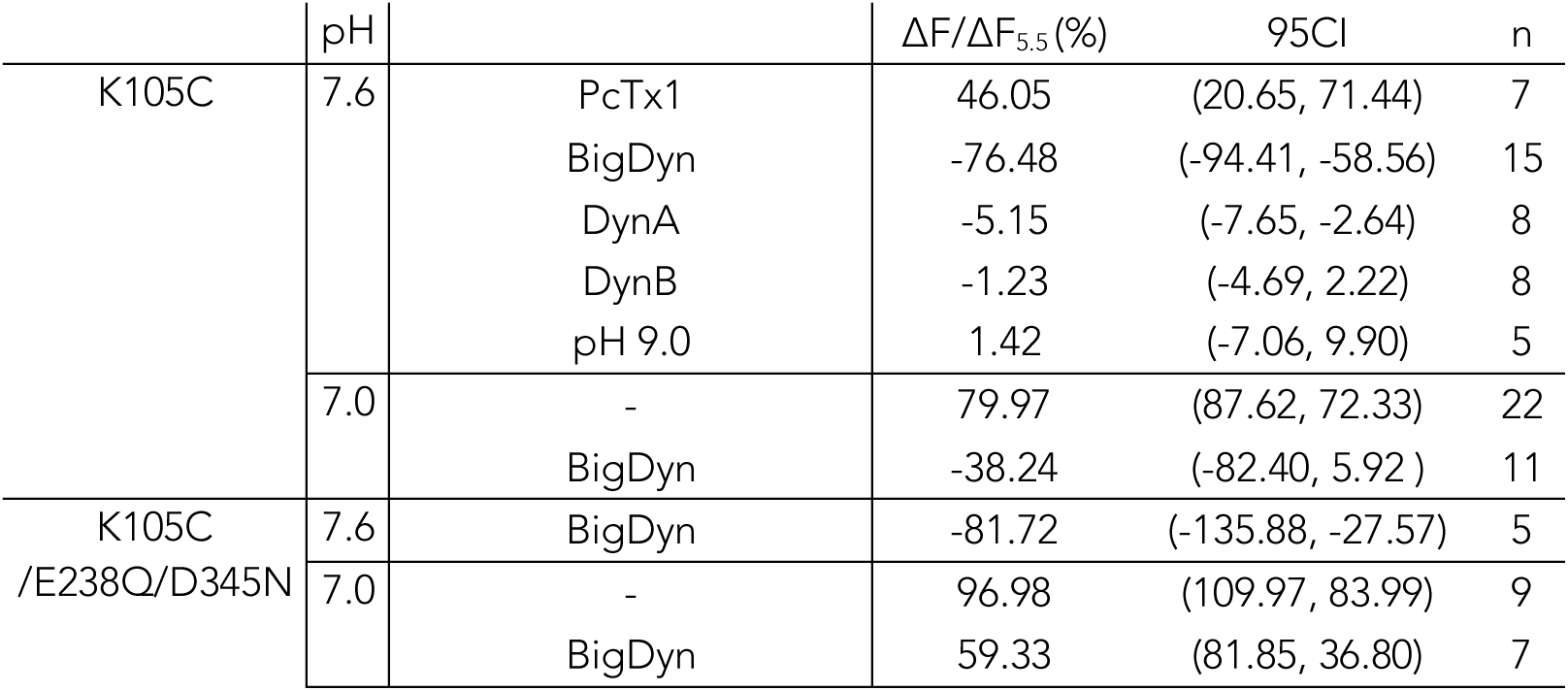
Peptide and pH-induced fluorescence changes at Alexa Fluor 488-labelled Lys105Cys. Mean fluorescence change (in % of that induced by pH 5.5 application, ΔF/ΔF_5.5_) induced by application of PcTx1 (0.3 µM), DynB (10 µM), DynA(10 µM), BigDyn (1 µM) or pH 9.0, with 95CI in parentheses; n = number of experiments.

| | pH | | $\Delta F/\Delta F_{5.5}$ (%) | 95CI | n |
| --- | --- | --- | --- | --- | --- |
| K105C | 7.6 | PcTx1 | 46.05 | (20.65, 71.44) | 7 |
|  |  | BigDyn | -76.48 | (-94.41, -58.56) | 15 |
|  |  | DynA | -5.15 | (-7.65, -2.64) | 8 |
|  |  | DynB | -1.23 | (-4.69, 2.22) | 8 |
|  |  | pH 9.0 | 1.42 | (-7.06, 9.90) | 5 |
|  | 7.0 | - | 79.97 | (87.62, 72.33) | 22 |
|  |  | BigDyn | -38.24 | (-82.40, 5.92 ) | 11 |
| K105C<br>/E238Q/D345N | 7.6 | BigDyn | -81.72 | (-135.88, -27.57) | 5 |
|  | 7.0 | - | 96.98 | (109.97, 83.99) | 9 |
|  |  | BigDyn | 59.33 | (81.85, 36.80) | 7 |

**Table S3:** pH-induced activation, steady-state desensitization and ΔF at Alexa Fluor 488-labelled Lys105Cys. pH_50_ and Hill coefficients for activation, steady-state desensitization and fluorescence change (ΔF/ΔF_pH5.5_), with 95CI stated in parentheses; n = number of experiments.

| | $pH_{50}$ | 95CI | Hill coefficient | 95CI | n |
| --- | --- | --- | --- | --- | --- |
| Activation | 6.61 | (6.52, 6.70) | 6.02 | (7.30, 4.74) | 5 |
| Steady-state desensitization | 7.06 | (7.03, 7.09) | 12.10 | (11.10, 13.10) | 5 |
| $\Delta F$ | 7.11 | (7.06, 7.20) | 8.49 | (12.24, 4.73) | 5 |

**Table S4:** Effect of alanine-substituted BigDyn analogs. Mean rescue of current after exposure to steady-state desensitization-inducing pH at WT mASIC1a, with 95CI in parentheses; the effect of the peptides compared to control was analyzed using a one-way ANOVA with a Dunnett’s multiple comparisons test, *, p < 0.05; **, p < 0.01; ***, p < 0.001; ****, p < 0.0001; n = number of experiments; n.s = not significant.

|  | Rescue (%) | 95CI (%) | n | p-value |  |
| --- | --- | --- | --- | --- | --- |
| Control | 4.03 | (1.81, 6.28) | 46 | - |  |
| Big Dyn | 81.82 | (76.39, 87.24) | 48 | <0.0001 | **** |
| Y1A | 30.24 | (14.90, 45.59) | 12 | 0.0003 | *** |
| R6A | 9.78 | (-2.07, 5.14) | 9 | 0.9989 | n.s |
| R7A | 4.97 | (1.16, 8.78) | 12 | 0.9999 | n.s |
| I8A | 60.84 | (44.23, 77.44) | 12 | <0.0001 | **** |
| R9A | 2.35 | (1.68, 3.01) | 11 | 0.9997 | n.s |
| K11A | 29.35 | (10.92, 47.77) | 9 | 0.0031 | ** |
| K13A | 57.56 | (41.24, 73.87) | 9 | <0.0001 | **** |
| W14A | 52.68 | (35.48, 69.87) | 10 | <0.0001 | **** |
| D15A | 88.44 | (79.83, 97.05) | 10 | <0.0001 | **** |
| K18A | 62.97 | (38.62, 87.31) | 7 | <0.0001 | **** |
| R19A | 42.73 | (31.10, 54.36) | 10 | <0.0001 | **** |
| R25A | 76.27 | (59.00, 93.54) | 10 | <0.0001 | **** |
| R26A | 47.86 | (20.45, 75.26) | 7 | <0.0001 | **** |
| K29A | 63.36 | (40.19, 86.52) | 8 | <0.0001 | **** |

**Table S5:** Effect of different dynorphins and truncated BigDyn analogs on mASIC1a steady-state desensitization. Mean rescue of current after exposure to steady-state desensitization-inducing pH at WT mASIC1a, with 95CI in parentheses; the effect of the peptides compared to control was analyzed using a one-way ANOVA with a Dunnett’s multiple comparisons test, *, p < 0.05; **, p < 0.01; ***, p < 0.001; **** p < 0.0001; n.s = not significant. n = number of experiments.

| Concentration | Peptide | Rescue | 95CI | n | p-value |  |
| --- | --- | --- | --- | --- | --- | --- |
| 1 μM | Control | 5.73 | (3.83, 7.63) | 68 | - | |
|  | BigDyn | 90.11 | (77.92, 102.30) | 12 | <0.0001 | **** |
|  | DynA | 8.36 | (5.49, 11.22) | 6 | 0.9995 | n.s. |
|  | DynB | 4.45 | (2.72, 6.19) | 6 | 0.9998 | n.s. |
|  | DynA + DynB | 5.57 | (2.14, 8.99) | 5 | >0.9999 | n.s. |
|  | Dyn(1-19) + DynB | 12.83 | (1.68, 23.98) | 6 | 0.9228 | n.s. |
|  | Dyn(2-32) | 14.46 | (6.25, 22.68) | 12 | 0.2669 | n.s. |
|  | Dyn(4-32) | 22.90 | (13.44, 32.37) | 13 | <0.0001 | **** |
|  | Dyn(6-32) | 63.80 | (49.46, 78.14) | 12 | <0.0001 | **** |
|  | Dyn(8-32) | 11.02 | (2.43, 19.62) | 12 | 0.915 | n.s. |
|  | Dyn(11-32) | 6.41 | (0.79, 12.03) | 18 | 0.9998 | n.s. |
|  | Dyn(14-32) | 3.52 | (0.48, 6.55) | 7 | 0.9996 | n.s. |
|  | Dyn(17-32) | 1.58 | (0.68, 2.49) | 13 | 0.9851 | n.s. |
| 3 μM | DynA | 31.02 | (8.45, 53.59) | 7 | <0.0001 | **** |
|  | DynB | 7.49 | (-2.27, 17.26) | 4 | 0.9997 | n.s. |

**Table S6:** Characterization pH dependence of steady-state desensitization of WT and mutant mASIC1a and WT hASIC1a. pH_50_ and Hill coefficients for steady-state desensitization (obtained with indicated activating pH), including resulting pH to induce steady-state desensitization. 95CI in parentheses; n = number of experiments, Glu373Gln/Glu411Gln was characterized on a single batch of oocytes.

|  | Steady-state<br>desensitization<br>pH<sub>50</sub> | 95CI | Hill<br>coefficient | 95CI | n | Activation<br>pH | Steady-state<br>densensitization<br>pH |
| --- | --- | --- | --- | --- | --- | --- | --- |
| WT | 7.24 | (7.23,<br>7.25) | 12.61 | (11.82,<br>13.40) | 64 | 5.6 | 7.1 |
| E182Q | 7.27 | (7.25,<br>7.29) | 14.04 | (12.83,<br>15.25) | 8 | 5.6 | 7.1 |
| E235Q | 7.08 | (7.06,<br>7.11) | 12.39 | (11.40,<br>13.38) | 8 | 5.6 | 6.9 |
| D237N | 7.07 | (7.05,<br>7.10) | 12.31 | (9.13,<br>15.49) | 8 | 5.6 | 6.9 |
| E238Q | 7.26 | (7.24,<br>7.28) | 13.87 | (8.87,<br>18.87) | 7 | 5.6 | 7.1 |
| E242Q | 7.14 | (7.07,<br>7.21) | 11.57 | (8.51,<br>14.63) | 6 | 5.6 | 6.9 |
| D259N | 7.14 | (7.13,<br>7.15) | 10.46 | (8.19,<br>12.73) | 6 | 5.6 | 7.0 |
| D345N | 7.24 | (7.19,<br>7.29) | 12.64 | (10.13,<br>15.15) | 9 | 5.6 | 7.1 |
| D349N | 7.38 | (7.35,<br>7.40) | 12.54 | (9.81,<br>15.28) | 11 | 5.6 | 7.2 |
| D237N/D345N | 7.06 | (7.04,<br>7.08) | 10.23 | (7.97,<br>12.49) | 14 | 4 | 6.9 |
| D237N/D349N | 7.10 | (7.07,<br>7.13) | 12.67 | (9.64,<br>15.70) | 5 | 5.6 | 6.95 |
| E238Q/D345N | 7.27 | (7.22,<br>7.32) | 9.61 | (9.00,<br>10.22) | 5 | 5.6 | 7.1 |
| E238Q/D349N | 7.33 | (7.31,<br>7.34) | 13.63 | (11.75,<br>15.50) | 8 | 5.6 | 7.2 |
| D253N/E254Q | 7.06 | (7.03,<br>7.09) | 15.93 | (13.09,<br>18.76) | 6 | 5.6 | 6.9 |
| E373Q/E411Q | 6.92 | (6.90,<br>6.93) | 19.39 | (9.48,<br>29.31) | 3 | 4 | 6.8 |
| hASIC1a WT | 7.07 | (7.03,<br>7.11) | 8.11 | (4.61,<br>11.61) | 6 | 5.6 | 6.8 |

**Table S7:** Effect of 1 µM BigDyn on steady-state desensitization of single and double mutant mASIC1a. Mean rescue of current after exposure to steady-state desensitization-inducing pH at WT and mutant mASIC1a in the absence (Control) and presence of 1 µM BigDyn (BigDyn), with 95CI in parentheses; the rescue in presence of BigDyn compared to Control was analyzed using an unpaired t-test, *, p < 0.05; **, p < 0.01; ***, p < 0.001; ****, p < 0.0001; n.s = not significant; n = number of experiments.

|  | Steady-state<br>desensitization<br>pH | Control |  |  | BigDyn |  |  | p-value |  |
| --- | --- | --- | --- | --- | --- | --- | --- | --- | --- |
|  |  | Rescue | 95CI | n | Rescue | 95CI | n |  |  |
| WT | 7.1 | 2.31 | (1.26, 3.36) | 45 | 82.81 | (77.80, 87.82) | 41 | <0.0001 | **** |
| E182Q | 7.1 | 0.93 | (-0.48, 2.33) | 7 | 82.01 | (65.55, 98.48) | 7 | <0.0001 | **** |
| E235Q | 6.9 | 3.30 | (0.90, 5.63) | 4 | 88.29 | (85.35, 91.22) | 6 | <0.0001 | **** |
| D237N | 6.9 | 3.95 | (2.07, 5.84) | 10 | 81.16 | (68.27, 94.05) | 14 | <0.0001 | **** |
| E238Q | 7.1 | 12.30 | (4.87, 19.74) | 9 | 69.74 | (50.55, 88.93) | 9 | <0.0001 | **** |
| E242Q | 6.9 | 4.64 | (-1.30, 10.57) | 6 | 92.86 | (82.27, 103.40) | 9 | <0.0001 | **** |
| D259N | 7.0 | 8.65 | (4.20, 13.10) | 9 | 69.59 | (56.38, 82.81) | 13 | <0.0001 | **** |
| D345N | 7.1 | 7.69 | (2.20, 13.17) | 8 | 58.92 | (42.53, 75.31) | 10 | <0.0001 | **** |
| D349N | 7.2 | 2.01 | (-1.49, 5.50) | 7 | 49.69 | (36.20, 63.19) | 12 | <0.0001 | **** |
| D237N/D345N | 6.9 | 2.24 | (0.10, 4.37) | 6 | 19.79 | (13.27, 26.32) | 10 | 0.0004 | *** |
|  | 6.95 | 2.27 | (0.56, 4.83) | 4 | 16.08 | (7.99, 24.18) | 10 | 0.0400 | * |
| D237N/D349N | 6.90 | 0.84 | (0.29, 13.98) | 5 | 5.53 | (-2.04, 13.09) | 4 | 0.0609 | n.s. |
|  | 6.95 | 2.63 | (0.72, 4.53) | 7 | 22.13 | (8.77, 35.48) | 8 | 0.0071 | ** |
| E238Q/D345N | 7.1 | 1.43 | (0.59, 2.27) | 10 | 6.766 | (1.69, 11.84) | 13 | 0.0606 | n.s. |
|  | 7.15 | 1.65 | (0.18, 3.13) | 6 | 13.62 | (7.30, 19.94) | 8 | 0.0026 | ** |
| E238Q/D349N | 7.1 | 0.70 | (0.24, 1.17) | 6 | 21.45 | (9.40, 33.51) | 8 | 0.0045 | ** |
|  | 7.2 | 2.79 | (0.39, 5.19) | 6 | 27.03 | (14.66, 39.40) | 10 | 0.0046 | ** |
| D253N/E254Q | 6.9 | 1.06 | (-0.02, 2.13) | 7 | 88.07 | (73.45, 102.70) | 11 | <0.0001 | **** |
| E373Q/E411Q | 6.8 | 2.25 | (-0.61, 5.11) | 5 | 141.1 | (91.66, 190.50) | 5 | <0.0001 | **** |

**Table S8:**
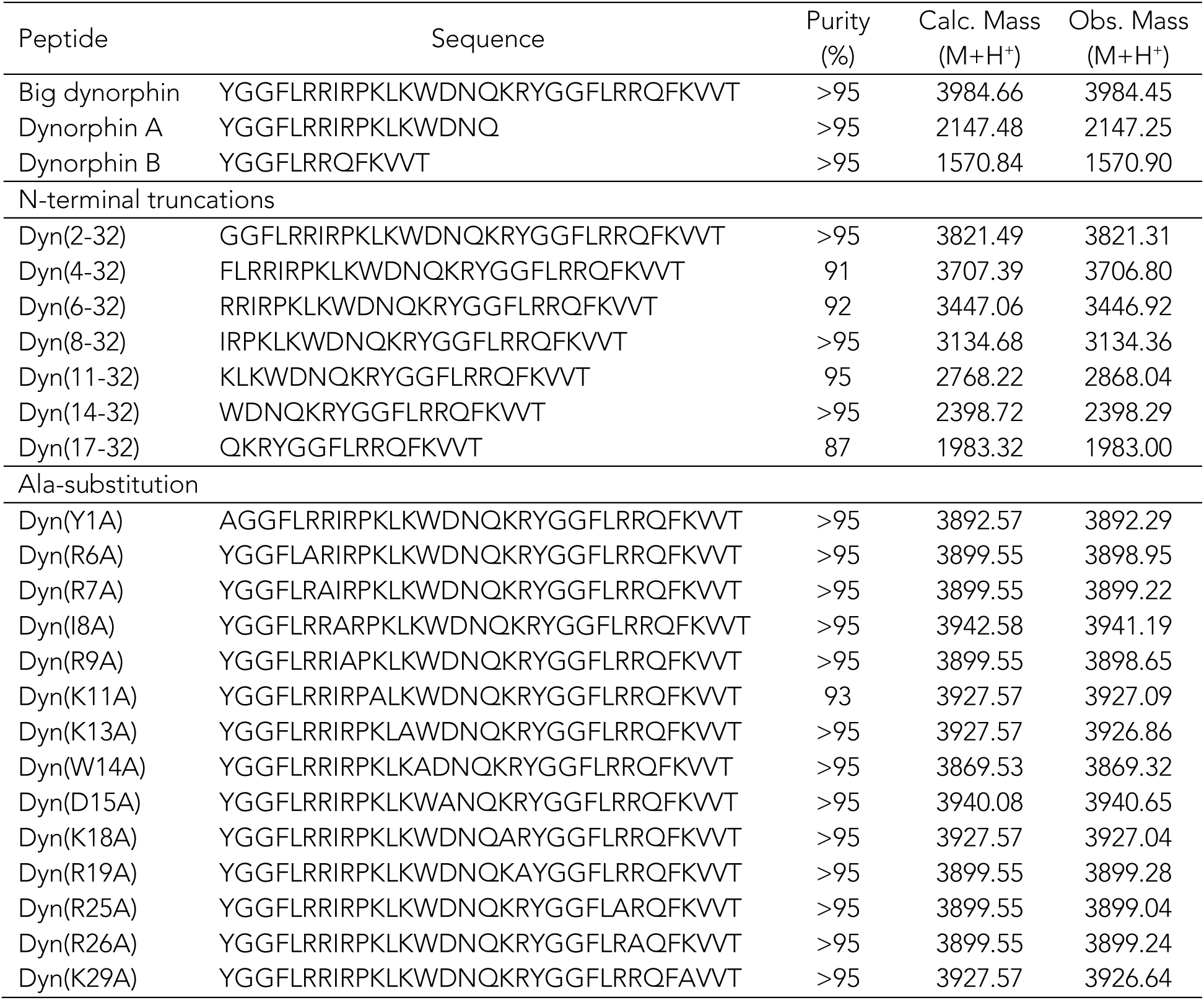
Synthesized peptides. Listed are peptide names, sequence, purity, as well as calculated and observed mass, respectively.

## Notes

#### Summary of Updates

Revised text and figures.

